# Transient excitability and synaptic consolidation support stable memory despite neural drift

**DOI:** 10.64898/2026.09.15.751773

**Authors:** Surbhit Wagle, Claudia Clopath

**Affiliations:** Bioengineering Department, Imperial College London

## Abstract

Long-term memories remain behaviorally stable despite turnover in the neuronal populations that encode them. This representational drift differs across brain regions: hippocampal representations can reconfigure within hours, whereas cortical ensembles remain comparatively stable over days to weeks. This asymmetry poses a challenge for systems consolidation, in which hippocampal activity is thought to instruct the formation of persistent cortical memory traces. We developed a two-region recurrent network model of the hippocampus (HPC) and anterior cingulate cortex (ACC) incorporating region-specific synaptic plasticity, excitability-dependent neuronal allocation, hippocampal-to-cortical coupling, and activity-dependent intrinsic plasticity. When the regions evolved independently, faster synaptic turnover in HPC produced pronounced drift, whereas persistent ACC connectivity preserved a more stable cortical ensemble. In the intact circuit, hippocampal input recruited ACC neurons but also propagated hippocampal variability into cortex, destabilizing the emerging cortical engram and impairing memory expression. A transient increase in the intrinsic excitability of recruited ACC neurons counteracted this instability by promoting repeated reactivation during an early consolidation window, stabilizing cortical ensemble membership without preventing hippocampal drift. Simulated erasure of learning-induced potentiation further reproduced the early dependence of memory on hippocampal, but not ACC, plasticity. Together, these results support a sequential mechanism in which hippocampal recruitment, transient cortical intrinsic plasticity, and persistent cortical synaptic plasticity transform a dynamic hippocampal representation into a stable cortical memory trace.

## Introduction

Long-term memories can persist for months or years, yet the neuronal populations that encode those memories are remarkably dynamic. Classical theories of memory assumed that stable behaviour arises from relatively stable memory engrams. However, advances in chronic neuronal recordings have challenged this view by demonstrating that the neurons participating in a representation can change substantially across days and weeks even when behaviour remains unchanged. This phenomenon, known as representational drift, has now been observed across numerous brain regions, including hippocampus [1–6], sensory cortex [7–12], parietal cortex [13] and motor cortex [14–16], suggesting that continual reorganization of neural representations is a fundamental property of the brain rather than an experimental artifact. Rather than viewing representational drift as a failure of memory storage, recent work has proposed that continual remodeling of neuronal ensembles may provide important computational advantages [17, 18]. Ensemble fluidity allows memories to incorporate new information while preserving existing knowledge, providing a mechanism for memory updating, generalization and behavioural flexibility. Slow fluctuations in intrinsic neuronal excitability, among others [10, 19–24], have emerged as one biological mechanism underlying representational drift [18, 25, 26]. Neurons experiencing transient increases in excitability are more likely to be recruited into active ensembles during learning and memory reactivation [27–29], leading to gradual turnover in engram membership over time. Such excitability-dependent allocation enables memories to evolve without requiring complete replacement of the underlying representation, potentially balancing stability with adaptability. These observations raise a fundamental challenge for theories of memory consolidation. According to the systems consolidation framework, newly acquired memories initially depend on hippocampal circuits before gradually becoming supported by distributed cortical networks through repeated offline reactivation and synaptic plasticity [30–33]. Extensive experimental evidence has demonstrated coordinated interactions between hippocampus and cortical regions such as the anterior cingulate cortex (ACC) during this process, with cortical representations becoming progressively stronger as hippocampal dependence decreases [34, 35]. Yet most computational theories of systems consolidation implicitly assume that hippocampal replay provides a relatively stable teaching signal that can be transferred to cortex over time. The existence of representational drift complicates this picture. If hippocampal engrams continuously reorganize because of excitability-driven changes in neuronal allocation, then cortical circuits are unlikely to receive an invariant teaching signal during consolidation. Instead, the information arriving in cortex is itself dynamic. Rule and colleagues proposed that drift may generate ongoing error signals between interconnected brain regions, allowing downstream circuits to continuously recalibrate their representations and preserve stable behaviour despite changing neural activity [36]. Likewise, theoretical studies have demonstrated that adaptive readout mechanisms can maintain stable decoding from unstable neural populations [37]. However, how such mechanisms interact with systems consolidation remains largely unexplored. In particular, it is unknown whether excitability-driven hippocampal drift can coexist with the formation of stable cortical engrams or whether additional biological mechanisms are required to prevent drift from propagating through memory networks. Recent experimental work suggests that intrinsic plasticity may provide such a stabilizing mechanism. Beyond transient excitability fluctuations that bias memory allocation, learning also induces activity-dependent increases in neuron-wide intrinsic excitability in neocortical engram neurons [38]. These changes persist over behaviourally relevant timescales and have been shown to regulate memory formation, allocation and precision [38]. Such findings suggest that intrinsic excitability may play a dual role in memory: promoting flexible recruitment of new neurons while simultaneously stabilizing previously established cortical ensembles during consolidation. Here, we develop a computational model to investigate how systems consolidation proceeds in the presence of ongoing representational drift. We constructed a two-region recurrent network consisting of hippocampal (HPC) and anterior cingulate cortex (ACC) populations with distinct synaptic plasticity timescales and excitability dynamics. We first show that differences in local plasticity naturally produce rapidly drifting hippocampal representations alongside relatively stable cortical engrams. We then demonstrate that, in a more biologically realistic network architecture, hippocampal drift propagates into cortex through feedforward interactions, challenging the stability of cortical memory representations. Finally, we show that activity-dependent intrinsic plasticity selectively stabilizes cortical engrams without preventing hippocampal drift, preserving behavioural output and reproducing experimentally observed temporal requirements for hippocampal and cortical plasticity [35]. Together, our results provide a mechanistic framework that reconciles representational drift with systems consolidation by identifying intrinsic plasticity as a key process enabling stable long-term memories to emerge from continuously evolving neural representations.

## Results

To investigate how stable behaviour can emerge despite ongoing reorganization of memory representations, we developed a two-region rate model with recurrent connections comprising a hippocampal (HPC) module and an anterior cingulate cortex (ACC) module (Fig. 1A). Each region consisted of excitatory neurons with recurrent all-to-all connectivity and were learned using Hebbian platicity rule. We simulated the network for 11 days protocol: the first day corresponded to an initial encoding of a memory and the other days corresponded to spontaneous or cue-induced reactivation of the neural ensamble (Fig. 1B, methods). Finally, intrinsic excitability was allowed to fluctuate across days, with distinct neuronal subpopulations exhibiting transient increases in excitability at different time points (Fig. 1C, methods).

**Figure 1:**
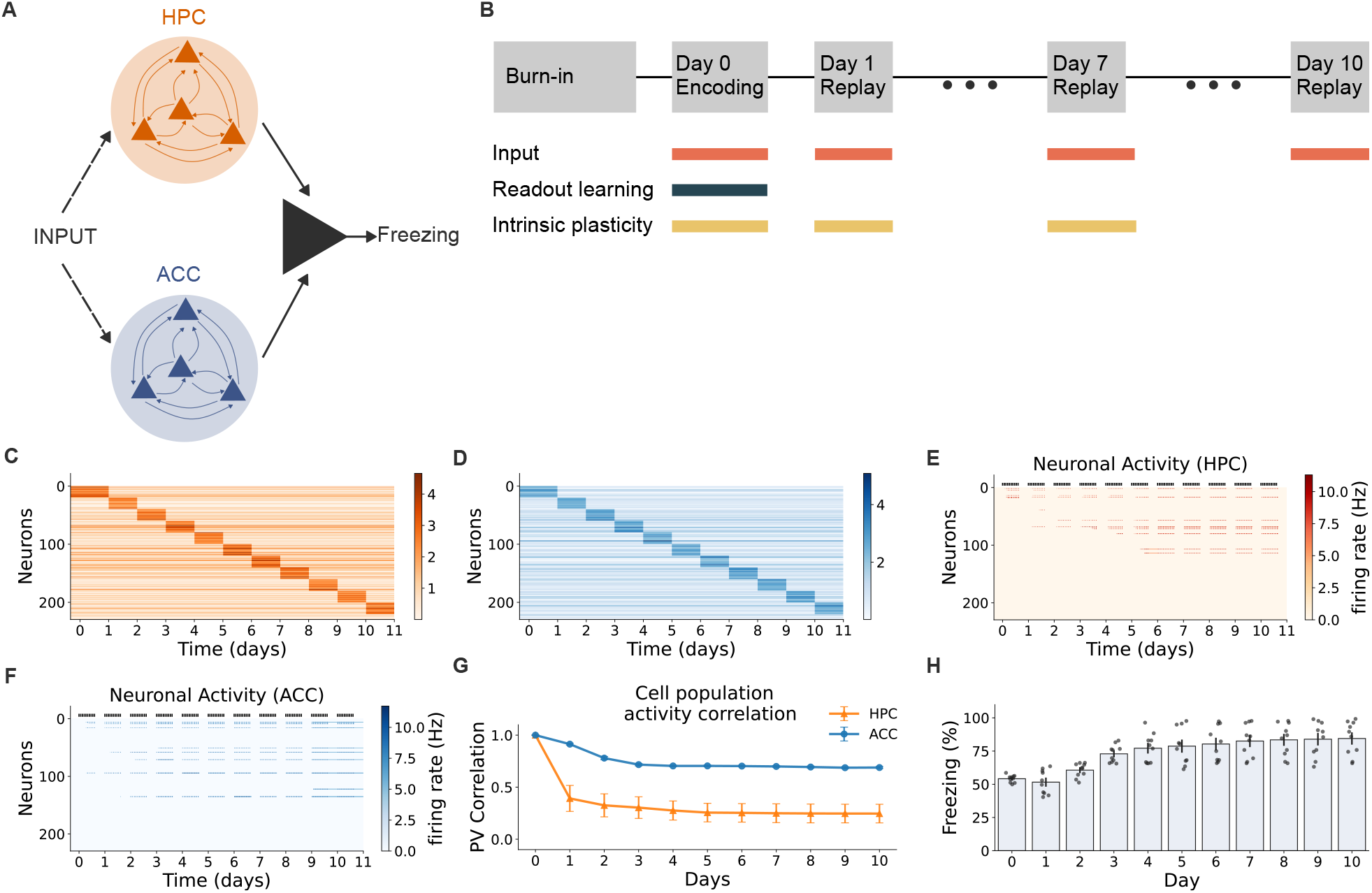
Synaptic plasticity timescale governs representational stability despite excitability turnover. **(A)** Conceptual framework. The model consists of a hippocampal (HPC) network and an anterior cingulate cortex (ACC) network, each composed of recurrently connected excitatory neurons. HPC and ACC differ in their plasticity timescales, with faster recurrent plasticity in HPC and slower recurrent plasticity in ACC. A behavioural performance neuron receives input from the memory representation and is trained during Encoding (FC). **(B)** Simulation protocol. Memories are encoded during FC and subsequently reactivated during offline consolidation epochs across multiple days. Representational drift emerges in HPC through excitability fluctuations and changing ensemble membership, whereas ACC representations gradually stabilize through recurrent strengthening and consolidation. **(C**,**D)** Distribution of intrinsic excitability (*ϵ*_*i*_) across neurons in **(C)** HPC network and **(D)** ACC network over time. During each stimulation epoch, a distinct subset of neurons exhibits elevated excitability (Methods). The model is used to study how stable behavioural output can be maintained despite ongoing engram reorganization in different circuits. **(E)** Activity of the HPC network across the simulation. Black tick marks above the heatmap indicate times when the external input was presented. As excitability is transferred to different neuronal groups across days, HPC activity progressively shifts toward newly excitable neurons. **(F)** Activity of the ACC network under the same input and excitability-turnover schedule. In contrast to HPC, ACC preserves a more stable active ensemble across days, while gradually incorporating additional neurons. **(G)** Population-vector correlation of HPC and ACC activity patterns relative to the day-0 encoding pattern. Correlations were computed from the mean activity vector for each day and averaged across simulations. HPC correlations decline rapidly, indicating high representational drift, whereas ACC correlations remain higher, indicating preservation of a stable engram core. **(H)** Freezing percentage across days, calculated from output-neuron firing rate (see Methods) and averaged over the 10 stimulus/replay repetitions of each day. Bars show mean*±*S.E.M. across simulations, with individual points representing single simulations. Freezing progressively approaches a high ceiling value, indicating that the downstream behavioral readout becomes saturated even while HPC and ACC population representations differ in their stability.

### Synaptic plasticity timescale governs stable engram cell allocation despite excitability turnover

To determine whether synaptic persistence can stabilize an engram during excitability turnover, first, we simulated independently driven HPC and CTX recurrent networks, without any connections between the two networks. In both regions, an excitability increase was changed to a new group of 20 neurons each day, continually changing which cells were intrinsically favored for recruitment (Fig. 1C&D). We then exposed both networks to a new memory by stimulating them with an external input. The two networks differed in their recurrent plasticity timescales. HPC synapses combined relatively rapid Hebbian potentiation with substantial decay, whereas CTX synapses learned more slowly and also had negligible decay rate.

HPC activity closely followed the changing excitability landscape (Fig. 1E). The population activated on each day progressively shifted toward newly excitable neurons, while recurrent connections formed within the currently recruited ensemble and earlier connections weakened. This distinction was also evident in the recurrent synaptic weights. In the HPC network, connections supporting previously active assemblies progressively weakened across days, while new connections strengthened among the neurons recruited by ongoing excitability turnover (Fig. S1A). In contrast, the ACC network maintained a stable core of recurrent weights associated with the original ensemble, while additional connections gradually strengthened as new neurons were incorporated into the cortical representation (Fig. S1B). Accordingly, mean population-vector (PV) correaltion, measured as the pearson correlation of average firing rate across the stimulation presentent time, to day-0 HPC representation decreased sharply to approximately 0.39 after one day and continued declining toward 0.25 by day 10 (Fig. 1G, orange line). These results indicate that the relatively short effective lifetime of HPC synaptic modifications allowed intrinsic excitability to repeatedly redirect engram allocation. ACC network on the other hand showed substantially greater representational stability despite undergoing the same excitability turnover (Fig. 1F,G blue line). The initial cortical ensemble remained active across later sessions, while newly excitable neurons were gradually incorporated into the representation. Consistent with this persistence, recurrent connections involving the original ensemble remained strong throughout the simulation. Day-0 population-vector correlation remained approximately 0.91 on day 1 and stabilized near 0.69 by day 10 (Fig. 1F,G blue line). Thus, ACC did not maintain a perfectly invariant population, but preserved a stable core representation while accommodating some recruitment of additional neurons. Together, these results show that excitability turnover does not necessarily produce equivalent engram turnover. Rapidly decaying recurrent plasticity caused HPC allocation to track the currently excitable population, whereas persistent CTX synapses retained the influence of earlier co-activation. The synaptic plasticity timescale therefore controls the balance between representational flexibility and stability: short-lived synaptic changes permit rapid engram reorganization, while persistent recurrent connectivity preserves a stable engram core despite continuing changes in intrinsic excitability.

To test how representational drift affects downstream behavioral expression, we simulated two additional output-readout conditions in which freezing was driven selectively by either the HPC or ACC network. In the HPC-output condition, only HPC-to-output plasticity was enabled, while ACC-to-output weights were set to zero. In this case, freezing was highest at post-encoding but decreased sharply after excitability turnover began, falling from approximately 46% on day 0 to approximately 27% by day 10 (Fig. S2A&B). This indicates that when behavior depends on the drifting HPC representation, the output readout becomes less consistently activated as the originally reinforced HPC ensemble is replaced by newly excitable neurons. In contrast, when the output was driven selectively by ACC, freezing progressively increased and approached a high ceiling value, rising from approximately 30% on day 0 to approximately 77% by day 10 (Fig. S2C&D). Thus, the more stable ACC representation provides a more reliable substrate for behavioral expression across days. Together, these readout-specific simulations suggest that representational drift is not only a population-level coding phenomenon, but can directly determine whether a downstream behavioral output is degraded or stabilized. One caveat of our model is that, in our model the neuronal activity in two networks evolved independetely of each other. Experimental work suggest that the coupled reactivation of neuronal assmeblies during sharp-wave ripples in the HPC and thalamo-cortical spindeles is necessary for consolidation [39–42]. In order, to study the effect of this coupled reactivation, we included a feed-forward weights between HPC and cortical network.

### In a realistic network architecture, hippocampal drift propagates to cortex

We next extended the model from two independent recurrent circuits to a coupled HPC–ACC architecture. In this model, HPC and ACC were each represented as recurrent rate networks, but ACC neurons additionally received, all-to-all feedforward excitatory connections from HPC (Fig. 2A&B). During encoding, the encoding memory activated both networks and trained the downstream output neuron. During subsequent reactivation sessions, HPC activity could drive ACC through the feedforward pathway, allowing co-active HPC and ACC ensembles to strengthen HPC→ACC synapses and ACC recurrent connections. We first examined whether this feedforward pathway was required for cortical recruitment. When HPC→ACC plasticity was removed (Fig. S3A), the HPC network continued to respond to the input and drifted across days as excitability shifted to newly recruited neurons (Fig. S3B). However, ACC activity was nearly absent, indicating that direct cortical input alone was insufficient to recruit a detectable ACC ensemble under these low-input conditions (Fig. S3C). This effect was aslo obervable in the recurrent synaptic weights, which matured and drifted for the HPC neuron, but failed to mature for the ACC network (Fig. S3D&E). Consistent with this failure of cortical recruitment, freezing behavior was strongly reduced after encoding and remained low across later days (Fig. S3F). Enabling HPC→ACC plasticity restored ACC recruitment during reactivation (Fig. 2B&C). In this condition, hippocampal activity provided an additional drive to ACC, allowing cortical neurons to become active and form a detectable ensemble. This increased behavioral output relative to the no-feedforward condition, indicating that HPC→ACC coupling supports downstream memory expression (Fig. 2E vs Fig.S3F, day 1 onwards). However, the recruited ACC representation remained only weakly correlated with the original day-0 pattern (Fig. 2D), and freezing did not return to encoding levels (Fig. 2E). Thus, hippocampal feedforward drive is necessary for engaging ACC in the memory network, but is not sufficient on its own to produce a stable cortical engram. This drift propagation was evident in quantitative measures of cortical ensemble similarity, which declined over time in the realistic model (Fig. 2C). Together, these simulations support HPC→ACC coupling as a recruitment mechanism during early systems consolidation. The hippocampal network can drive cortical participation and support behavioral expression, but additional cortical stabilization mechanisms are required to preserve the ACC engram and behavior over time.

**Figure 2:**
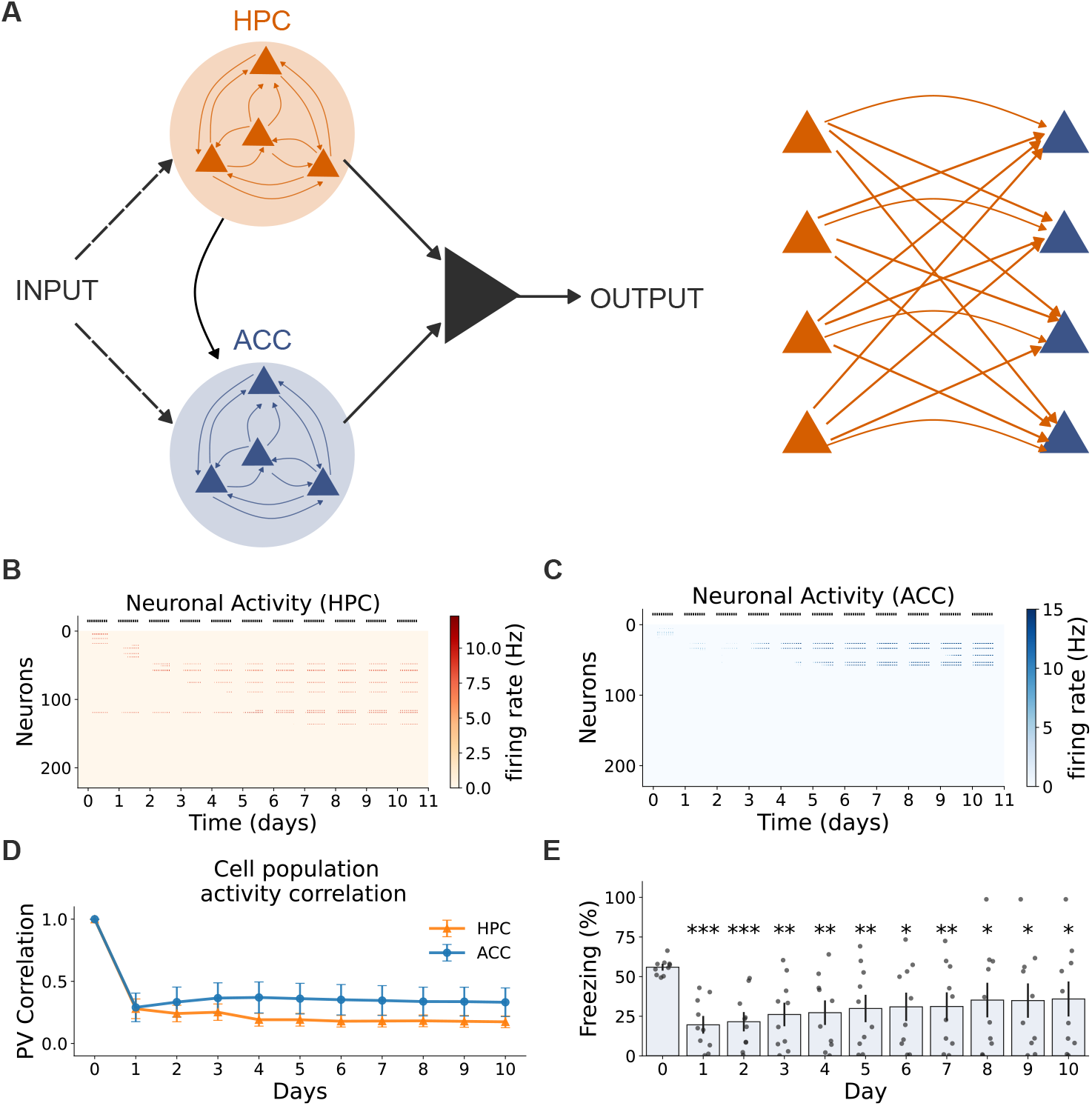
A more realistic network architecture reveals that hippocampal drift can propagate to cortical representations. **(A)** Schematic of the extended two-region model with all-to-all inter-regional (shown on right) connectivity between HPC and ACC. In addition, the behavior is only decoded from the ACC neurons. In contrast to the simplified model, this architecture allows ongoing hippocampal activity to more strongly influence cortical dynamics during consolidation and recall. **(B, C)** Example activity patterns in HPC and ACC across consolidation days. Although HPC engrams continue to exhibit representational drift, the denser inter-regional coupling causes this instability to spread to ACC, resulting in progressive turnover of cortical ensemble membership. **(D)** Quantification of ensemble similarity in HPC and ACC across days, measured as population vector correlation. Compared with the simplified model, ACC representations are less stable and show increased drift over time. **(E)** Consequences for behavioural performance. Because cortical representations are no longer fully stable, decoding performance becomes more variable across consolidation, indicating that synaptic consolidation alone is insufficient to fully buffer behavioural output against hippocampal drift in the realistic model. These results identify drift propagation from HPC to ACC as a key challenge for consolidation in recurrent multi-region networks.

Thus, although recurrent cortical plasticity still promoted engram formation, it was no longer sufficient on its own to fully stabilize the ACC representation against inherited hippocampal variability. Consequently, behavioural performance also became less stable than in the simplified model (Fig. 2D). These results reveal an important limitation of synaptic consolidation alone: in distributed recurrent circuits, hippocampal drift can spread to cortex and compromise the stability of the consolidated memory trace. This observation motivated us to test whether an additional stabilizing mechanism, implemented here as activity-dependent intrinsic plasticity, could prevent drift propagation and preserve cortical engram stability.

### Activity-dependent excitability modulation stabilizes cortical engrams

After establishing that HPC→ACC coupling is required to recruit ACC activity, we next asked whether an additional cortical stabilization mechanism was needed to preserve the recruited ACC ensemble. This was motivated by recent experimental work showing that contextual fear learning induces a transient increase in the intrinsic excitability of nascent ACC engram neurons [38]. This increase is expressed during the early phase of systems consolidation and largely returns to baseline within approximately one week. Moreover, manipulating the excitability state of tagged ACC neurons specifically during this early window altered their subsequent maturation, remote reactivation, memory strength, contextual precision, and susceptibility to interference. These findings suggest that intrinsic excitability plasticity does not itself constitute a permanent memory-storage mechanism, but instead provides a transient permissive signal that biases which recently recruited cortical neurons remain eligible for repeated reactivation and subsequent stabilization.

In line with these findings, ACC neurons that became active during memory reactivation received a transient increase in intrinsic excitability in our model. This excitability boost decayed over approximately 7 days, approximating the restricted temporal window of intrinsic excitability plasticity observed experimentally (Fig. 3A). The excitability dynamics of all ACC neurons showed across the population, different neurons became transiently excitable on successive days, producing a continuous daily turnover in the excitable neuronal cohort (Fig. 3D). Within this dynamically changing population, neurons belonging to the developing engram, highlighted by red stars, underwent intrinsic plasticity following their recruitment and retained elevated excitability across subsequent days. Although these neurons also showed a gradual decay toward baseline, their transient excitability increased their responsiveness to hippocampal and recurrent cortical input and enhanced their probability of being recruited again during later reactivation events. Thus, intrinsic plasticity did not eliminate population turnover, but biased the competition created by this turnover in favor of previously recruited engram neurons. Rather than allowing each day’s newly excitable cohort to replace the existing representation, previously recruited neurons remained preferentially eligible for repeated reactivation, thereby maintaining their participation in the developing cortical engram.

**Figure 3:**
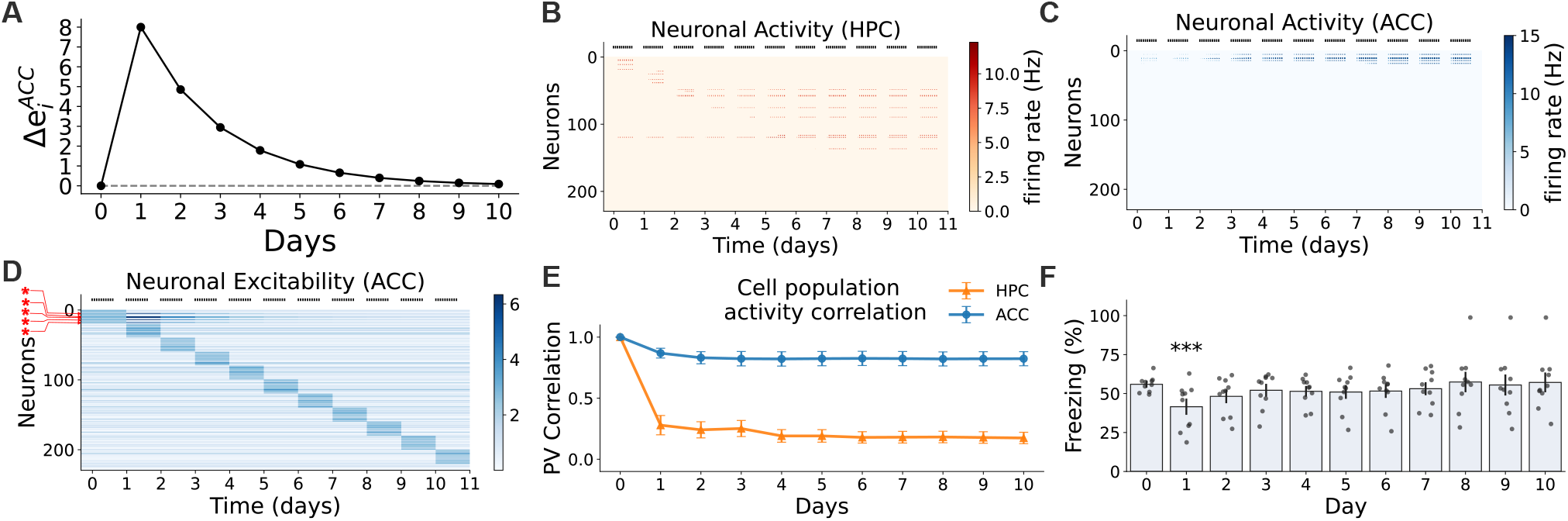
ACC intrinsic plasticity stabilizes the cortical engram after HPC-driven recruitment. **(A)** Temporal dynamics of intrinsic-plasticity mechanism. ACC neurons activated during reactivation receive a transient excitability boost that decays over days. **(B)** HPC activity across days with ACC intrinsic plasticity. HPC activity continues to drift across neuronal cohorts, indicating that the manipulation does not prevent hippocampal turnover. **(C)** ACC activity across days with intrinsic plasticity. Previously recruited ACC neurons remain preferentially active, producing a stable cortical ensemble despite ongoing excitability turnover. **(D)** Dynamics of neuronal excitability in ACC neurons. Engram neurons that fire above the threshold of 5 Hzs undergo IP of excitability (shown in red stars on left). **(E)** Population-vector correlation relative to the day-0 encoding pattern. HPC correlation declines rapidly, whereas ACC correlation remains high across days when intrinsic plasticity is present. **(F)** Freezing behavior averaged over the last 10 presentations of each day. Intrinsic plasticity restores behavioral output to near encoding levels, in contrast to the no-intrinsic-plasticity condition.

Adding ACC intrinsic plasticity strongly stabilized the cortical representation (Fig. 3C and E). In the nointrinsic-plasticity condition, ACC population-vector correlation with the day-0 encoding pattern remained low, reaching only approximately 0.22 by day 10 (Fig. 2D). With intrinsic plasticity, ACC correlation remained high across the simulation, staying near 0.83 by day 10 despite continued drift in the HPC network (Fig. 3E). Importantly, HPC activity was essentially unchanged by the ACC intrinsic-plasticity manipulation: HPC correlation still declined to approximately 0.13 by day 10 (Fig. 3B and E). This indicates that the stabilization was specific to the cortical network and was not caused by preventing hippocampal drift. Intrinsic plasticity also restored behavioral expression. Without ACC intrinsic plasticity, freezing fell from approximately 57% on day 0 to approximately 28% by day 10 (Fig. 2E). With intrinsic plasticity, freezing recovered to approximately 60% by day 10, returning close to the original encoding level (Fig. 3F). Thus, transient intrinsic excitability modulation converted a weakly recruited and unstable ACC ensemble into a persistent cortical memory trace capable of supporting behavioral output.

Together, these simulations suggest a two-step consolidation mechanism. First, HPC→ACC coupling recruits ACC neurons into the memory network during reactivation. Second, transient intrinsic plasticity maintains the recruited neurons in a consolidation-competent state, increasing their likelihood of participating in subsequent reactivation events. Repeated recruitment can then support the local synaptic changes required for long-term stabilization after the excitability boost has decayed. HPC remains a drifting network, but its early recruitment of ACC is sufficient to seed a cortical ensemble whose membership is initially protected by intrinsic plasticity and subsequently preserved by local cortical synaptic plasticity. This interpretation is consistent with the proposal of Hadzibegovic et al. that early intrinsic excitability plasticity acts as a permissive tag governing the maturation and eventual fate of nascent ACC engram neurons, rather than serving as the enduring physical substrate of the memory itself [38].

### ACC intrinsic plasticity plays a causal role in establishing stable cortical engrams

Having established that transient intrinsic plasticity can stabilize HPC-recruited ACC neurons despite ongoing daily excitability turnover, we next asked when during consolidation this mechanism is required. For this, we then tested whether intrinsic plasticity was required throughout consolidation or only during a restricted temporal window. To address this, we blocked the tagged-cell excitability boost during selected consolidation days while leaving the daily excitability turnover of the wider neuronal population intact. Blocking intrinsic plasticity during the early consolidation window, on days 1–2, largely abolished ACC stabilization (Fig. 4A&C). ACC day-10 correlation fell to approximately 0.21, closely matching the condition in which intrinsic plasticity was absent throughout the simulation (Fig. 4E). Freezing was similarly reduced, reaching only approximately 28% by day 10 (Fig. S4A).

**Figure 4:**
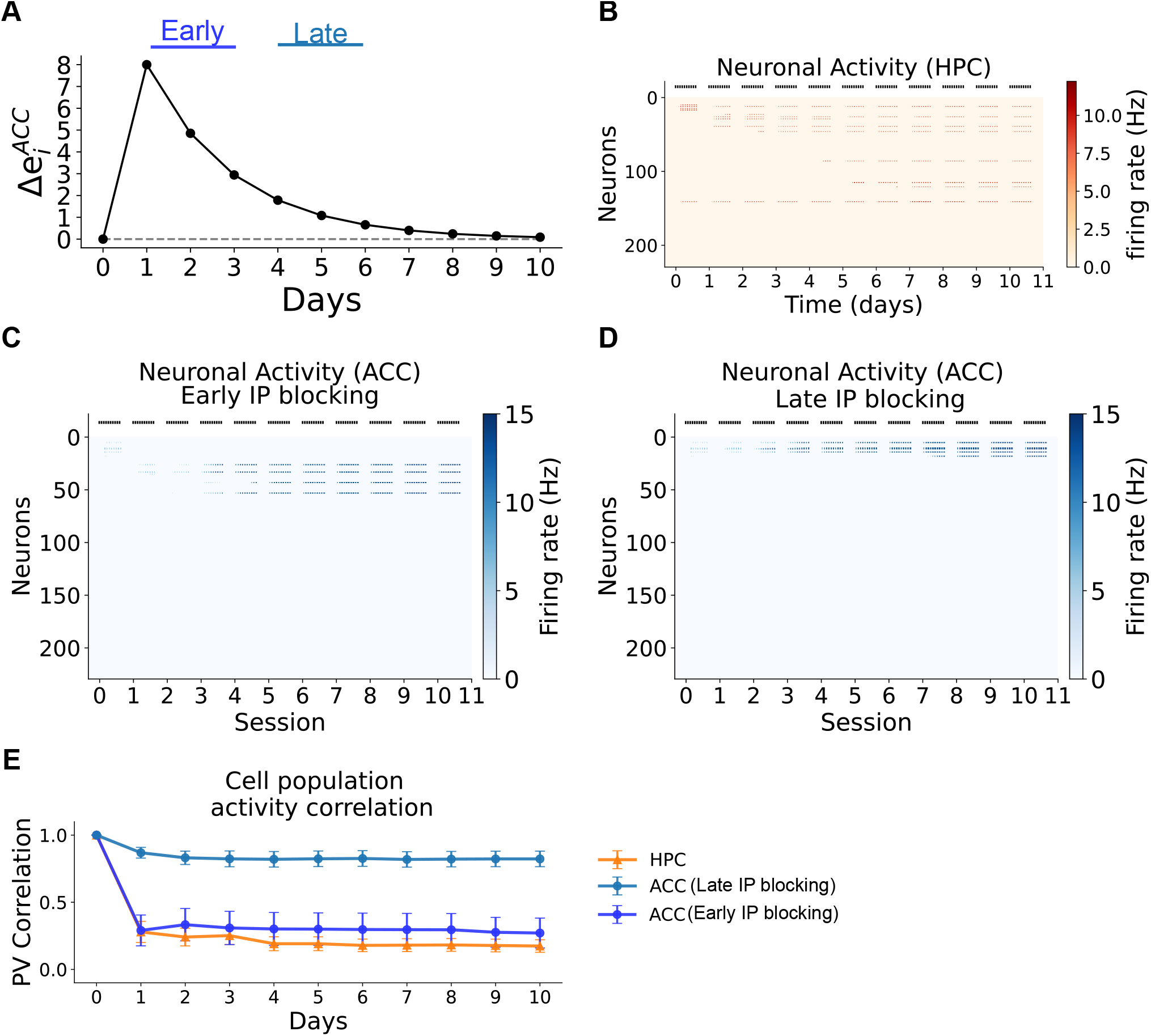
ACC intrinsic plasticity is causally required for stable cortical engram formation. **(A)** ACC neurons activated during reactivation receive a transient excitability boost that decays over days. We implmented *in-silico* blocking of the intrensic platicity (early, day 1-2, and late day 4-5 after encoding). (B) HPC activity across days with ACC intrinsic plasticity. HPC activity continues to drift across neuronal cohorts, indicating that the manipulation does not prevent hippocampal turnover. **(C)** ACC activity across days with intrinsic plasticity blcoked during the early days (day 1-2). Without the early intrensic plasticity, the engram neurons in ACC do not maintain stable firing and new engram neurons gets recruited. **(D)** ACC activity across days with intrinsic plasticity blcoked during the late days (day 4-5). Previously recruited ACC neurons remain preferentially active, producing a stable cortical ensemble. **(E)** Population-vector correlation relative to the day-0 encoding pattern. HPC correlation declines rapidly.PV correaltion in ACC neurons also declines sharply when the intrinsic plasticity is blocked at early time, whereas a late block doesn’t affect PV correaltion and representation stability.

In contrast, blocking intrinsic plasticity during later periods, on days 4–5, had little effect on the final cortical representation or behavioral output. ACC day-10 correlation remained near 0.75, while day-10 freezing remained near 56% (Fig. 4D&E, Fig. S4B). Thus, once the recruited cortical ensemble had been repeatedly reactivated and stabilized, later interruptions of the excitability boost had comparatively weak consequences. This temporal dissociation indicates that intrinsic plasticity is particularly important during an early consolidation window, when nascent ACC engram neurons must compete with the continuously changing population of neurons favored by daily excitability turnover.

Together, these simulations refine the proposed two-step mechanism of systems consolidation. HPC→ACC coupling initially recruits cortical neurons during memory reactivation, but recruitment alone is insufficient to ensure their long-term participation in the memory ensemble. During the early consolidation period, intrinsic plasticity maintains recently recruited ACC neurons in a temporarily privileged, consolidation-competent state, increasing their likelihood of being reactivated despite ongoing neuronal turnover. Repeated reactivation can then support the local cortical synaptic changes required to establish a persistent engram. Once this representation has been stabilized, continued intrinsic excitability enhancement becomes less critical for subsequent memory expression.

### The model reproduces the qualitative behavioral effect of temporally targeted plasticity erasure

Although intrinsic plasticity can maintain recently recruited ACC neurons in a consolidation-competent state, a persistent cortical engram ultimately requires stabilization of the synaptic connections linking these neurons. This idea is supported by Goto et al., who directly examined the spatiotemporal requirement for long-term potentiation during systems memory consolidation [35]. Hence, we asked whether our model could reproduce the behavioral consequences of experimentally blocking plasticity during memory consolidation. In the experiments of Goto et al., recently induced LTP was erased using an optogenetic approach based on a fusion of cofilin with the photosensitizer SuperNova (CFL–SN) [35]. Cofilin accumulates in dendritic spines undergoing structural potentiation, allowing illumination-induced chromophore-assisted light inactivation to disrupt the actin reorganization supporting recently established LTP within a restricted temporal window. CFL–SN was expressed in either hippocampal CA1 or the ACC, and light was delivered at selected post-learning time points to determine when recently induced synaptic potentiation in each region was necessary for memory consolidation [35]. In the model, LTP erasure was implemented by comparing synaptic weights before and after learning, identifying the most strongly potentiated synapses from the distribution of positive weight changes, and selectively removing their learning-induced component while preserving baseline connectivity (see Methods). Under full erasure, the selected synapses were reset to their pre-learning values without silencing the corresponding neurons or altering non-potentiated connections. Memory was assessed experimentally using an inhibitory-avoidance task in which mice learned to avoid crossing from an illuminated compartment into a dark compartment where they had previously received a foot shock. Experimental memory expression was quantified as the training-induced change in crossover latency, with a larger increase in the time taken to enter the dark compartment indicating stronger avoidance memory. In the model, memory expression was instead quantified as freezing-like output activity. Because crossover latency and freezing-like activity have different units and arise from different behavioral assays, we compared the direction and timing of the effects of LTP erasure rather than directly equating their absolute magnitudes.

First, we examined manipulations targeting hippocampal plasticity. In the model, erasing potentiated synaptic changes in the hippocampal network either immediately after encoding or during the first offline consolidation period reduced subsequent freezing relative to control simulations (Fig. 5A,C, right). This result was consistent with the experimental findings of Goto et al., who erased recently induced LTP in hippocampal CA1 either 2 min after inhibitory-avoidance training or during the first post-learning sleep period [35]. Both manipulations impaired memory measured on the following day, as indicated by a smaller training-induced increase in crossover latency than in control mice (Fig. 5A,C, left). In contrast, erasing recently induced CA1 LTP more than one day after learning did not significantly affect subsequent crossover latency (Fig. 5E,left), indicating that the critical period for hippocampal potentiation was restricted to the early post-learning phase. The model reproduced this temporal dependence: erasing hippocampal synaptic changes one day after encoding did not substantially reduce freezing relative to the corresponding control condition (Fig. 5E, right). Thus, despite the difference in behavioral readout, the model captured both the early sensitivity and the later resistance to hippocampal LTP erasure observed experimentally. These results suggest that hippocampal plasticity makes an essential contribution immediately after learning and during the first offline consolidation period, but becomes less vulnerable to disruption once these early plasticity events have occurred.

**Figure 5:**
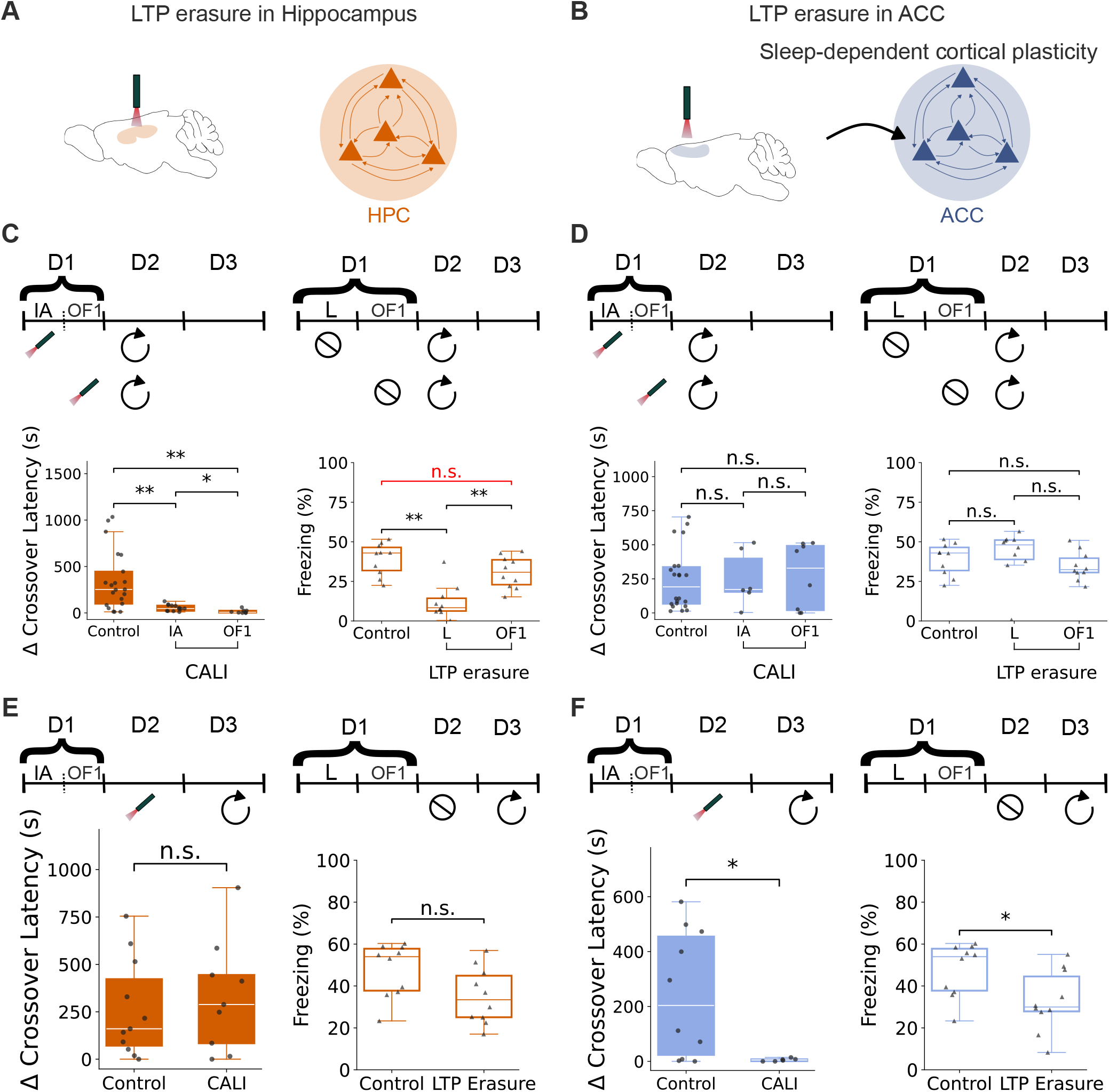
The model reproduces the qualitative behavioral consequences of temporally targeted plasticity blockade. **(A)** *Left:* Schematic of experimental hippocampal LTP erasure using chromophore-assisted light inactivation (CALI) in dorsal CA1. *Right:* Schematic of the HPC network in which potentiated recurrent synaptic weights were reset to model LTP erasure (see Methods).**(B)** *Left:* Schematic of experimental cortical LTP erasure using CALI in the ACC. *Right:* Schematic of the ACC network in which potentiated recurrent and HPC→ACC feedforward synaptic weights were reset to model LTP erasure (see Methods). **(C)** *Top left:* Experimental timeline showing hippocampal LTP erasure using CALI either 2 min after learning or during the first offline period on day 1. Memory was subsequently tested on day 2. *Top right:* Corresponding model timeline in which potentiated HPC recurrent weights were reset either immediately after encoding or after the first offline reactivation session, followed by recall testing on day 2. *Bottom left:* Hippocampal LTP erasure 2 min after learning or during the first offline period produced a smaller training-induced increase in crossover latency than in control mice, indicating impaired inhibitory-avoidance memory (Control: N =21, CALI: N(IA) = 13, CALI N(OF1) N=8. Contro;l VS CALI IA: U=227, corrected p=2.8e-3 (holm), CALI IA vs CALI OF1: U=87, corrected p=1.2e-1 (holm), Control vs CALI OF1: U=158, corrected p=1e-3 (holm)). *Bottom right:* The model reproduced this qualitative effect, with HPC weight resetting immediately after encoding or after the first offline reactivation resulting in significantly lower freezing than in control simulations without weight resetting (N=10 simulation for each condition, Control vs LTP Erasure L: U=96, corrected p=1.7e-3 (holm), LTP Erasure L vs LTP Erasure OF1: U=9, corrected p=2.2e-3 (holm), Control vs LTP Erasure OF1 : U=74, corrected p=7.5e-2 (holm)). **(D)** Same arrangement as in **(C)**, but with LTP erasure targeted to the ACC. *Top left:* Experimental timeline showing ACC LTP erasure either 2 min after learning or during the first offline period on day 1. *Top right:* Corresponding model timeline in which potentiated ACC recurrent and HPC→ACC feedforward weights were reset at the analogous time points. *Bottom left:* Neither early ACC manipulation significantly altered the training-induced change in crossover latency relative to control mice (Control: N=22, CALI: N(IA) = 6, N(OF1) = 8, Control vs CALI IA: U=63, corrected p=1.00 (holm), CALI IA vs CALI OF1: U=24, corrected p=1.00 (holm), Control vs CALI OF1 : U=90, corrected p=1.00 (holm)). *Bottom right:* Consistently, ACC weight resetting at the corresponding time points did not significantly alter freezing relative to control simulations (N=10 simulation for each condition, Control vs LTP erasure L: U=35, corrected p=0.54 (holm), LTP erasure L vs LTP erasure OF1: U=77, corrected p=0.13 (holm), Control vs LTP erasure OF1: U=65, corrected p=0.54 (holm)). **(E)** *Top left:* Experimental timeline showing hippocampal LTP erasure using CALI one day after learning, followed by inhibitory-avoidance testing on day 3. *Top right:* Corresponding model timeline in which potentiated HPC recurrent weights were reset at the analogous delayed time point. *Bottom left:* Delayed hippocampal LTP erasure did not significantly affect memory, as shown by a change in crossover latency comparable to that of control mice (Control: N=11, CALI: N=9, U=44.5, p=0.7323). *Bottom right:* Similarly, delayed HPC weight resetting did not substantially alter freezing relative to control simulations (N=10 simulation for each condition, Control vs LTP Erasure: U=76, p=0.0539). **(F)** *Top left:* Experimental timeline showing ACC LTP erasure during sleep on day 2, one day after learning. *Top right:* Corresponding model timeline in which potentiated ACC recurrent and HPC→ACC feedforward weights were reset at the analogous delayed consolidation time point. *Bottom left:* Erasure of recently induced ACC LTP during sleep significantly reduced the training-induced increase in crossover latency, indicating impaired inhibitory-avoidance memory (Control: N=10, CALI: N =10, U=49.5, p=4e-2). *Bottom right:* In agreement to the experimental result, delayed ACC weight resetting substantially reduce freezing in the model (N =10 simulations for each condition, U=81, p=2.1e-2), indicating that the current Hebbian plasticity rule does not reproduce the delayed, sleep-dependent requirement for rapid ACC potentiation.

We then examined analogous manipulations in the cortical/ACC component of the model. Experimentally, erasing ACC LTP either 2 min after learning or during the first offline period did not significantly affect subsequent recall, as measured by the training-induced change in crossover latency (Fig. 5B,D, left). The model reproduced this early resistance to ACC LTP erasure: resetting potentiated ACC synaptic weights immediately after encoding or during the corresponding first offline period did not substantially reduce freezing relative to control simulations (Fig. 5B,D, right). Thus, both the experimental and modelling results indicate that behaviorally critical ACC potentiation is not yet established during the earliest post-learning period.

Goto et al. next erased recently induced ACC LTP during sleep on the day after learning. In contrast to the earlier manipulations, mice receiving LTP erasure during this delayed offline period exhibited a significantly smaller increase in crossover latency at the subsequent memory test, indicating impaired inhibitory-avoidance memory [35]. Importantly, applying the same manipulation during wakefulness did not produce a comparable impairment, indicating that the requirement for ACC LTP was both delayed and brain-state dependent. A conventional Hebbian implementation with a constant ACC learning rate was unable to reproduce this temporal dissociation (Fig. S6A-C). Under this condition, cortical potentiation accumulated gradually across successive reactivation sessions, and disrupting plasticity during a single delayed offline period did not significantly reduce subsequent freezing. Because learning was distributed across consolidation periods, potentiation established before or after the targeted intervention could compensate for the disrupted episode. Thus, temporally homogeneous Hebbian plasticity captured gradual cortical strengthening but was insufficient to generate the delayed, step-like dependence on ACC LTP observed experimentally.

We therefore introduced a state- and time-dependent modulation of cortical plasticity. ACC recurrent plasticity remained weak during encoding and the earliest consolidation period but increased during NREM-like reactivation from day 2 onward. This modification retained the same underlying Hebbian learning rule while concentrating cortical potentiation within delayed offline consolidation periods. With this mechanism, erasing potentiated ACC synapses during the delayed offline period produced a corresponding reduction in freezing relative to control simulations (Fig. 5F). The updated model therefore reproduced the qualitative temporal dissociation observed by Goto et al.: early ACC LTP erasure had little effect, whereas disruption of potentiation during a later offline consolidation window impaired subsequent memory expression. Further support for a flexible cortical consolidation window comes from Liu et al. [43]. Experimentally, erasing recently induced ACC LTP on either day 2 or day 3 alone did not significantly impair subsequent inhibitory-avoidance memory, whereas erasing LTP during the offline periods on both days produced a significant reduction in crossover latency. These findings suggest that ACC consolidation is not tied to a single obligatory post-learning day. Instead, either of several available offline periods can support the required cortical potentiation, provided that at least one remains intact.

The constant-learning-rate model also failed to reproduce this cumulative effect: ACC potentiation was distributed across multiple reactivation sessions, allowing synaptic strengthening outside the targeted periods to maintain behavioral output even when consecutive consolidation sessions were disrupted. In contrast, the updated model reproduced the qualitative experimental pattern (Fig. S5). Resetting ACC potentiation during either day 2 or day 3 alone produced little effect on subsequent freezing, consistent with cortical strengthening occurring during the remaining intact high-plasticity period. Erasing potentiation across both days, however, removed the available high-plasticity consolidation opportunities and significantly reduced freezing relative to control simulations.

Together, these comparisons indicate that a conventional constant-rate Hebbian rule is insufficient to account for the temporal organization of ACC plasticity revealed by LTP-erasure experiments. Introducing a delayed increase in Hebbian learning specifically during NREM-like reactivation was sufficient to produce both the stepwise transition toward cortical dependence observed by Goto et al. and the flexible, compensatory consolidation window reported by Liu et al. The results therefore suggest that the critical feature is not a fundamentally different cortical learning rule, but temporal gating of the efficacy of Hebbian plasticity during offline consolidation.

## Discussion

Our results support a multistage account of systems consolidation in which hippocampal recruitment, transient intrinsic excitability, and persistent cortical synaptic plasticity serve complementary functions. Differences in recurrent plasticity timescales allowed hippocampal representations to remain flexible while cortical representations were comparatively stable. However, once the networks were coupled, hippocampal input was neccesary to recruit ACC neurons. However, the hippocampal input also transmitted hippocampal variability to ACC circuit. Activity-dependent intrinsic plasticity stabilized the recruited ACC population by temporarily preserving its competitive advantage, allowing repeated reactivation to establish persistent cortical connectivity. These findings suggest that stable cortical memory cannot be explained by slow synaptic decay alone. Hippocampal reactivation can provide a teaching signal for cortical recruitment, but a changing hippocampal population may also continually alter the pattern of cortical input. A local cortical allocation mechanism is therefore required to ensure that previously recruited neurons remain responsive while the cortical engram matures.

Transient intrinsic plasticity fulfilled this stabilizing role in the model. Rather than eliminating daily excitability turnover, it biased competition in favor of recently recruited ACC neurons. This interpretation is consistent with Hadzibegovic et al., who showed that nascent ACC engram neurons undergo an early and transient increase in intrinsic excitability that influences their later maturation and contribution to remote memory [38]. Intrinsic plasticity may therefore act as a permissive consolidation signal rather than as a permanent memory substrate. Importantly, the same transient excitability state may also create a period of vulnerability to interference. Hadzibegovic et al. showed that exposure to an interfering experience during the early consolidation window impaired subsequent remote-memory expression, whereas the same interference outside this window had little effect [38]. Chemogenetic suppression of the original ACC engram during the interfering experience reduced its recruitment into the competing representation and protected the original memory. Thus, elevated excitability may have two opposing consequences: it can promote repeated reactivation and maturation of the original engram, but it can also increase the probability that the same neurons are allocated to temporally adjacent experiences. Intrinsic plasticity may therefore define a transient period in which cortical engrams are both consolidation-competent and particularly susceptible to interference.

The temporal specificity of the intrinsic-plasticity manipulation further suggests that this mechanism is most important while cortical engram membership remains vulnerable. By repeatedly favoring the same neurons, elevated excitability increases their probability of coactivation and creates the conditions under which local recurrent synapses can subsequently strengthen. Once this connectivity is established, the ensemble becomes less dependent on continued excitability enhancement. Intrinsic and synaptic plasticity can therefore be viewed as complementary stages of stabilization: intrinsic plasticity preserves which neurons remain eligible for repeated recruitment, whereas synaptic plasticity converts this repeated coactivation into a persistent network representation.

The LTP-erasure results further support this temporal organization. The model reproduced the early requirement for hippocampal plasticity reported by Goto et al., as well as the relative resistance of memory to ACC LTP erasure immediately after learning or during the earliest offline period [35]. A conventional hebbian implementation with a constant ACC learning rate, however, was insufficient to reproduce the delayed sensitivity to cortical LTP erasure observed experimentally. Under this condition, ACC potentiation accumulated gradually across reactivation sessions, such that synaptic changes formed outside a targeted intervention period could compensate for erasure. Introducing a delayed, NREM-dependent increase in the ACC learning rate resolved this discrepancy. In the updated model, cortical plasticity remained weak during encoding and early consolidation but became substantially stronger during later NREM-like reactivation. This temporal gating concentrated ACC potentiation within delayed offline periods while retaining the same underlying Hebbian learning rule. With this modification, erasing ACC potentiation during the delayed consolidation window reduced subsequent freezing, reproducing the qualitative effect reported by Goto et al. [35]. Thus, the experimental temporal dissociation did not require replacing Hebbian learning with a fundamentally different rule, but did require its efficacy to vary across consolidation states and time.

The updated model also reproduced the flexible consolidation window reported by Liu et al. [43]. Experimentally, erasing ACC LTP during either day 2 or day 3 alone spared subsequent memory, whereas erasure across both offline periods impaired recall. A constant-learning-rate model could not reproduce this cumulative effect because cortical potentiation was distributed across many reactivation sessions. In contrast, when strong ACC plasticity was restricted to delayed NREM-like periods, disruption of either single window could be compensated for by potentiation during the remaining intact period, whereas erasure across both windows substantially reduced memory expression. These findings suggest that stepwise cortical consolidation can emerge from temporally gated Hebbian plasticity in which at least one intact high-plasticity offline period is sufficient for rapid ACC stabilization. The state-dependent increase in ACC plasticity implemented here is consistent with experimental evidence that NREM sleep provides a cortical state particularly favorable for synaptic modification. Spindle-related firing patterns can induce neocortical LTP, while cortical spindles are associated with synchronized dendritic *Ca*^2+^ activity and neuronal co-firing within the temporal window required for spike-timing-dependent plasticity [44–46]. Moreover, experimentally enhancing the coordination of hippocampal ripples with cortical slow oscillations and spindles promotes prefrontal network reorganization and memory consolidation, whereas disrupting appropriately timed NREM interactions impairs consolidation [33, 47, 48]. Thus, the transient increase in the ACC Hebbian learning rate during NREM-like reactivation should be interpreted as a simplified representation of the enhanced opportunity for synaptic modification created by sleep-specific oscillatory and dendritic dynamics.

The experimental and modelling erasure procedures nevertheless remain mechanistically distinct. CFL– SN selectively targets recently induced structural LTP within a restricted temporal interval, whereas the model resets positive learning-related changes in selected ACC recurrent and HPC→ACC synapses while preserving baseline connectivity and allowing later repotentiation. The updated model therefore captures the temporal organization of the behavioral effects more closely, but does not explicitly represent the molecular induction, stabilization, or persistence of LTP. Future extensions could distinguish these phases more explicitly and incorporate physiological signals associated with NREM sleep, including hippocampal replay and thalamocortical oscillations, as mechanisms that gate the transient increase in cortical plasticity.

The model additionally provides a potential link between intrinsic plasticity and silent cortical engrams. Silent engrams retain memory-related information but are not effectively recruited by natural cues, although direct artificial activation can restore memory expression [31, 49, 50]. Prefrontal engram neurons may be allocated early after learning but remain immature and naturally inaccessible until they undergo hippocampal-dependent maturation [30, 51, 52]. Systems consolidation may therefore involve not only the formation of a cortical ensemble, but also its transition from a silent to an accessible state. Transient intrinsic plasticity may facilitate this transition by maintaining the responsiveness of newly recruited ACC neurons while their recurrent connections mature. Without this excitability advantage, an ACC trace may be initially allocated but inconsistently reactivated, leaving it insufficiently connected to support natural cue-driven recall. Repeated reactivation could progressively strengthen engram-to-engram synapses until cortical pattern completion becomes sufficient to activate the memory independently of elevated intrinsic excitability. This interpretation generates several experimental predictions. Blocking intrinsic plasticity during early consolidation should reduce natural remote recall but may leave an ACC trace that can still be revealed by direct optogenetic or chemogenetic stimulation of learning-tagged neurons. Rescue of behavior would indicate that the manipulation produced a silent or inaccessible engram rather than complete memory loss. Early intrinsic-plasticity blockade should also reduce reactivation overlap across consolidation sessions and weaken the progressive strengthening of synapses between ACC engram neurons [34]. The timing of excitability restoration should also be important. Enhancing the excitability of tagged ACC neurons during early consolidation should promote repeated recruitment, cortical synaptic maturation, and lasting natural recall. In contrast, increasing excitability only during remote testing may transiently recover behavior without restoring stable accessibility on later tests. This distinction would separate temporary retrieval rescue from genuine engram maturation. LTP-erasure experiments provide a complementary test of this framework. Direct activation of ACC engram neurons after delayed or consecutive LTP erasure could determine whether the manipulation leaves behind a silent cortical trace. Behavioral rescue would suggest that synaptic erasure primarily prevents natural cue-driven pattern completion. Failure to rescue would indicate that cortical LTP is required to maintain the functional memory substrate itself. Combining early intrinsic-plasticity blockade with delayed LTP erasure could further test the proposed sequence in which intrinsic excitability maintains ensemble eligibility and synaptic potentiation establishes persistent connectivity.

Several limitations should be considered. HPC and ACC were represented as simplified rate-based recurrent networks without explicit hippocampal subfields, inhibitory populations, dendritic mechanisms, neuromodulation, or structured anatomical connectivity. Reactivation was imposed in discrete sessions, and sleep states were not modelled explicitly. Excitability turnover and intrinsic plasticity were implemented phenomenologically, and behavioral expression was represented by a simplified freezing-like output. The model should therefore be interpreted as identifying computational principles rather than reproducing the full biological circuitry of systems consolidation.

Overall, the results suggest that hippocampal input initially recruits a cortical memory representation, transient intrinsic plasticity preserves the recruited cells during an early competitive period, and persistent synaptic plasticity converts repeated reactivation into stable recurrent connectivity. This framework helps explain how a changing hippocampal population can seed a comparatively stable cortical engram and suggests that intrinsic plasticity may help transform an initially silent cortical trace into a mature and naturally accessible memory.

## Acknowledgements

We thank the members of the Clopath Lab for their valuable input and fruitful discussions. This work was supported by Wellcome Trust 200790/Z/16/Z, EPSRC EP/R035806/1, ERC MotorAdapt 101169605.

## Author contributions

S.W. and C.C. designed research; S.W. performed research; S.W. and C.C. wrote the manuscript.

## Conflict of interest

The authors declare no competing interests.

## Methods

### Network architecture

We implemented a multi-region recurrent rate-based network model to capture interactions between hippocampal (HPC; medial temporal lobe) and cortical anterior cingulate cortex (ACC) populations during memory encoding, replay, and consolidation. The model consisted of two recurrent neural populations, one hippocampal and one cortical, together with a single output neuron that served as a behavioral readout of conditioned freezing.

The hippocampal and cortical populations each contained *N* units, with firing-rate vectors

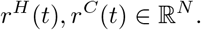

The output neuron had scalar activity *r*^*O*^(*t*). The hippocampal and cortical populations received external inputs, recurrent inputs, and activity-dependent global inhibition. The cortical population additionally received feedforward input from the hippocampal population. The output neuron received input from the hippocampal and cortical populations and was used to quantify freezing behavior.

### Rate dynamics

Neural activity evolved according to first-order rate dynamics:

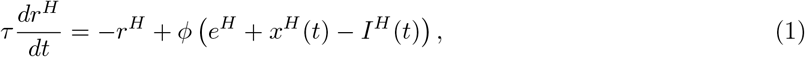

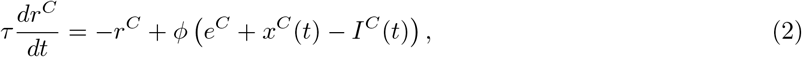

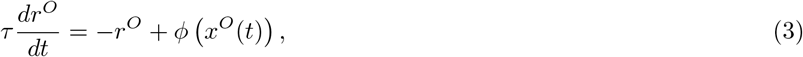

where *τ* = 20 is the neural time constant and

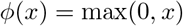

is a rectified linear activation function. The vectors *e*^*H*^ and *e*^*C*^ denote the intrinsic excitability of hippocampal and cortical units, respectively.

### Synaptic inputs

The total synaptic input to each population was defined as:

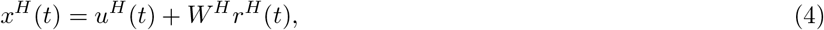

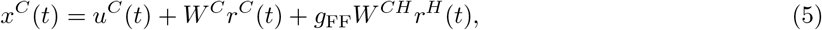

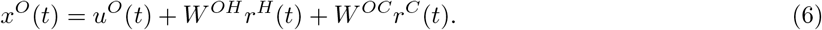

Here, *u*^*H*^ (*t*), *u*^*C*^(*t*), and *u*^*O*^(*t*) are external inputs to the hippocampal, cortical, and output populations, respectively. The matrices *W* ^*H*^ and *W* ^*C*^ denote recurrent hippocampal and cortical connectivity. The matrix *W* ^*CH*^ denotes feedforward hippocampus-to-ACC projections, and *g*_FF_ controls the effective strength of hippocampo-cortical feedforward coupling. The matrices *W* ^*OH*^ and *W* ^*OC*^ denote output weights from the hippocampal and cortical populations to the output neuron, respectively.

The output neuron received a mild guiding signal, *u*^*O*^(*t*), during encoding. This signal was set to 1 on day 0 and was set to 0 during all subsequent simulation days.

The parameter *g*_FF_ acted as a state-dependent gate on hippocampo-cortical communication. When *g*_FF_ = 1, hippocampal activity was transmitted to ACC through the learned feedforward weights. When *g*_FF_ = 0, this feedforward drive was functionally silenced without modifying the underlying synaptic weights.

### Global inhibition

Both hippocampal and cortical populations were subject to activity-dependent global inhibition:

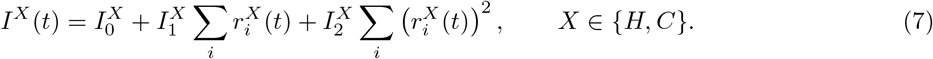

This inhibition provided population-level stabilization by suppressing activity as a function of the total and squared population firing rate. The parameters *I*_0_, *I*_1_ and *I*_2_ are reported in Table 1.

**Table 1:** Parameters used for simulating the different model circuits.

| Parameter | Fig. 1 | Fig. 2 | Fig. 3,4 | Fig. 5 |
| --- | --- | --- | --- | --- |
| $N_{\text{HPC}}$ | 230 | 230 | 230 | 230 |
| $N_{\text{ACC}}$ | 230 | 230 | 230 | 230 |
| $N_{\text{days}}$ | 11 | 11 | 11 | 11 |
| $N_{\text{rep}}$ | 10 | 10 | 10 | 10 |
| $T_{\text{stim}}$ | 100 | 100 | 100 | 100 |
| $T_{\text{IR}}$ | 100 | 100 | 100 | 100 |
| $T_{\text{ID}}$ | 1000 | 1000 | 1000 | 1000 |
| $I_{\text{cue}}$ | 13 | 13 | 13 | 13 |
| $E_{\text{HPC}}$ | 1.6 | 1.7 | 1.7 | 1.7 |
| $E_{\text{ACC}}$ | 1.6 | 1.7 | 1.7 | 1.7 |
| $E_{\text{mod}}$ | — | — | 8 | 8 |
| $\eta_{\text{HPC}}$ | $1.25 \times 10^{-3}$ | $1.25 \times 10^{-3}$ | $1.25 \times 10^{-3}$ | $1.25 \times 10^{-3}$ |
| $\lambda_{\text{HPC}}$ | $1 \times 10^{-3}$ | $1 \times 10^{-3}$ | $1 \times 10^{-3}$ | $1 \times 10^{-3}$ |
| $\eta_{\text{ACC}}$ | $5 \times 10^{-4}$ | $1 \times 10^{-5}$ | $1 \times 10^{-5}$ | $3 \times 10^{-6}$ to $3 \times 10^{-5}$ (see methods) |
| $\lambda_{\text{ACC}}$ | $1 \times 10^{-8}$ | $1 \times 10^{-8}$ | $1 \times 10^{-8}$ | $1 \times 10^{-8}$ |
| $\eta_{\text{HPC} \rightarrow \text{ACC}}$ | — | $1 \times 10^{-3}$ | $1 \times 10^{-3}$ | $1 \times 10^{-3}$ |
| $\lambda_{\text{HPC} \rightarrow \text{ACC}}$ | — | $1 \times 10^{-6}$ | $1 \times 10^{-6}$ | $1 \times 10^{-6}$ |
| $\eta_{\text{HPC} \rightarrow \text{OP}}$ | $1 \times 10^{-3}$ | $1 \times 10^{-3}$ | $1 \times 10^{-3}$ | $1 \times 10^{-3}$ |
| $\eta_{\text{ACC} \rightarrow \text{OP}}$ | $2 \times 10^{-3}$ | $2 \times 10^{-3}$ | $2 \times 10^{-3}$ | $2 \times 10^{-3}$ |
| $I_0, I_1, I_2$ | 9, 0.5, 0.05 | 9, 0.5, 0.05 | 9, 0.5, 0.05 | 9, 0.5, 0.05 |
| $N_{\text{sim}}$ | 10 | 10 | 10 | 10 |

### Intrinsic excitability fluctuations and intrinsic plasticity

Each unit was assigned an intrinsic excitability value that modulated its responsiveness to synaptic input. Baseline excitabilities were drawn independently from a half-normal distribution:

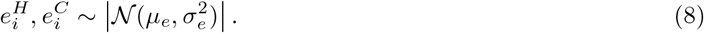

To model learning-dependent intrinsic plasticity, we introduced a transient increase in intrinsic excitability following encoding. Neurons that were active during encoding were marked as tagged units. For the ACC population, the set of tagged neurons was defined as

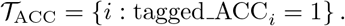

At the beginning of each simulated day *d*, cortical excitability was reset to its baseline value:

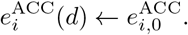

For tagged ACC neurons, excitability was then increased during a temporally restricted intrinsic-plasticity window:

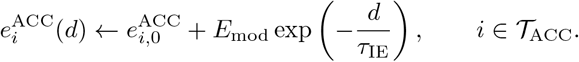

Here, *E*_mod_ is the magnitude of the excitability boost and *τ*_IE_ is the decay time constant of the intrinsic-excitability modulation. This formulation captures a transient post-learning increase in intrinsic excitability. Such increases have been proposed to act as permissive or tagging mechanisms that bias neurons toward future plasticity and stabilize memory representations. In particular, recent experimental work has shown that learning induces a time-limited increase in intrinsic excitability in neocortical engram neurons, which regulates memory formation, allocation, and precision [38]. Consistent with this framework, intrinsic plasticity in our model defines a temporal window during which tagged neurons are more likely to participate in subsequent plasticity and network reorganization. See Table 1 for specific values of parameters used to generate different figures.

### Activity tagging

During simulation, neurons were tagged when their activity exceeded a threshold *θ*_tag_:

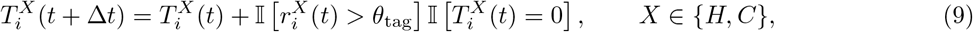

where I(·) is the indicator function. Thus, once a neuron crossed the tagging threshold, it remained tagged for subsequent intrinsic-plasticity updates.

### Simulation protocol

Each simulation began with a burn-in period of 1000 time steps without external input, allowing network activity to stabilize before learning. The network was then simulated across multiple days. Each day comprised 10 stimulation or reactivation epochs of 100 time steps, separated by 100-time-step inter-repetition intervals without external input. These repeated epochs were followed by a 1000-time-step period without external input.

During encoding on Day 0, external cue input was presented using a smooth temporal ramp, and readout learning was enabled by setting *s*_*O*_ = 1. Subsequent days represented offline sleep sessions. During each offline session, the cue was presented during the repeated NREM-like reactivation epochs, using the same temporal ramp as during encoding, but readout learning was disabled by setting *s*_*O*_ = 0. The intervening periods without external input represented inter-repetition rest intervals, and the extended period following the reactivation epochs represented a REM-like phase.

Recall sessions were performed on predetermined days by presenting the cue with all synaptic plasticity disabled. The synaptic weights and neuronal states present immediately before recall were restored after each recall test, ensuring that recall did not modify subsequent network consolidation.

### Synaptic plasticity

Recurrent synapses within the hippocampal and cortical populations, as well as feedforward synapses from the hippocampus to the ACC, evolved according to thresholded Hebbian plasticity with synaptic weight decay. The thresholded activity of neuron *i* in population *X* was defined as

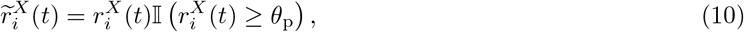

where *X* ∈ *{H, C}* denotes the hippocampal or cortical population, respectively, I(·) is the indicator function, and *θ*_p_ = 0.2 is the plasticity threshold.

Recurrent weights within each population were updated according to

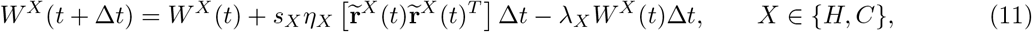

where *η*_*X*_ and *λ*_*X*_ denote the learning and decay rates, respectively, and *s*_*X*_ controls whether plasticity is enabled in population *X*.

Feedforward hippocampus-to-ACC weights were updated according to

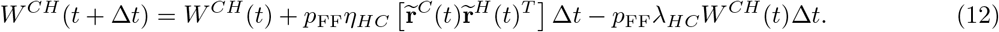

Thus, *p*_FF_ = 1 enabled hippocampus-to-ACC plasticity, whereas *p*_FF_ = 0 froze the feedforward weights. Following each update, synaptic weights were constrained to the interval [0, 1]. The plasticity parameters are reported in Table 1.

### Sleep-dependent cortical plasticity

Sleep-dependent cortical plasticity was implemented by modulating the learning rate of recurrent ACC synapses during offline reactivation. The baseline cortical learning rate was *η*_*C*_ = 3 *×* 10^−6^ during encoding, the first offline session, inter-repetition intervals, and REM-like periods. From offline Day 2 onward, the cortical learning rate was increased tenfold to

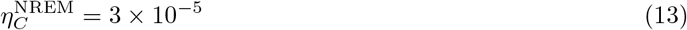

specifically during the NREM-like reactivation epochs. At the end of each reactivation epoch, the learning rate returned to its baseline value. Consequently, enhanced cortical synaptic plasticity was temporally restricted to reactivation during later offline sessions, providing a model of delayed, sleep-dependent strengthening of recurrent cortical connections.

### Readout learning

The output neuron provided a scalar readout of conditioned freezing. During encoding, readout weights were updated using a supervised signal *s*_*O*_(*t*). The weights onto output neuron were updated according to

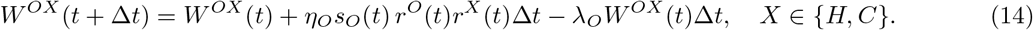

Readout learning was enabled on day 0 and disabled on subsequent simulation days. Refer to Table 1 for specific values of plasticity parameters.

### Freezing quantification

The output neuron was interpreted as a behavioral readout of conditioned freezing. To express output activity in behavioral units, we converted the output-neuron firing rate into percent freezing using a linear saturating transformation. If *r*_out_(*t*) is the firing rate of the output neuron at time *t*, freezing was defined as

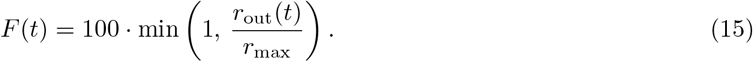

Here, *F* (*t*) is the predicted freezing percentage and *r*_max_ is the output firing rate corresponding to 100% freezing. In all simulations, we set

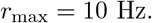

Thus, output rates at or above 10 Hz were mapped to 100% freezing, whereas zero output activity was mapped to 0% freezing. For each simulated day, average freezing was quantified from all the 10 the stimulus-presentation epochs.

### Simulation of LTP erasure

LTP erasure was implemented as a synapse-specific reversal of positive weight changes relative to a reference weight matrix. For each plastic projection, a reference matrix *W* ^ref^ was stored immediately before the onset of the manipulation. Following each stimulation or replay event on an LTP-erasure day, the change in each synaptic weight was calculated as

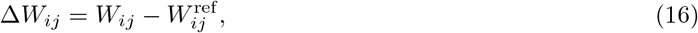

where *W*_*ij*_ is the current weight from presynaptic neuron *j* to postsynaptic neuron *i*. Synapses that had undergone potentiation (Δ*W*_*ij*_ *>* 0) were returned to their reference values:

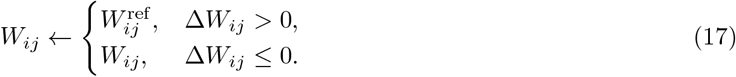

The resulting weights were constrained to the interval *W*_*ij*_ ∈ [0, 1]. Thus, the manipulation selectively removed newly acquired potentiation while preserving unchanged synapses and synaptic depression.

For hippocampal LTP erasure, this operation was applied to the recurrent hippocampal weight matrix. For cortical LTP erasure, it was applied to both the recurrent cortical weight matrix and the feedforward hippocampal-to-cortical weight matrix. Network activity and synaptic plasticity evolved normally during each stimulation or replay event, after which the positive weight changes were reversed. Consequently, the manipulation prevented the persistence of LTP-like changes without directly altering neuronal activity, intrinsic excitability, or synapses that had not potentiated.

For consecutive LTP-erasure days, the same pre-manipulation reference matrix was retained so that potentiation accumulating across the complete intervention period was removed relative to a common baseline. On non-erasure days, the reference matrix was updated to the current weights. The number of erased synapses was recorded as the number of elements satisfying Δ*W*_*ij*_ *>* 0.

### Statistical analysis

Statistical analyses were performed in Python using SciPy. Data are shown as boxplots representing the median and interquartile range, with individual simulation runs or animals overlaid as separate points. Pairwise differences between specified groups were assessed using two-sided Mann–Whitney *U* tests. When more than one pairwise comparison was performed within an analysis, *p*-values were adjusted using the Holm step-down procedure to control the family-wise error rate. For a single pre-specified comparison, the unadjusted *p*-value was reported.

## Digitisation of published experimental data

Experimental data used for the comparisons in Figure 5, figure S5 and S6 were obtained by digitising the relevant graphical results reported by Goto et al. (2021) [35] and Liu et al., (2025) [43]. Individual data points were extracted from the published figures using [WebPlotDigitizer](https://automeris.io/) and were replotted using custome Pyton scripts for the present study. These values represent secondary estimates derived from the published graphs rather than original raw experimental data. Consequently, small differences from the underlying measurements may arise from image resolution and digitisation precision. Sample sizes and details of the experimental interventions were taken from the original publication. No new animal experiments were performed for these comparisons.

## Code Availability

The code used for the generating the simulations are available online: [GitHub Repository](https://github.com/surbhit21/Systems_consolidation.git)

## Supplmental Information

### Supplementary figures

**Figure S1:**
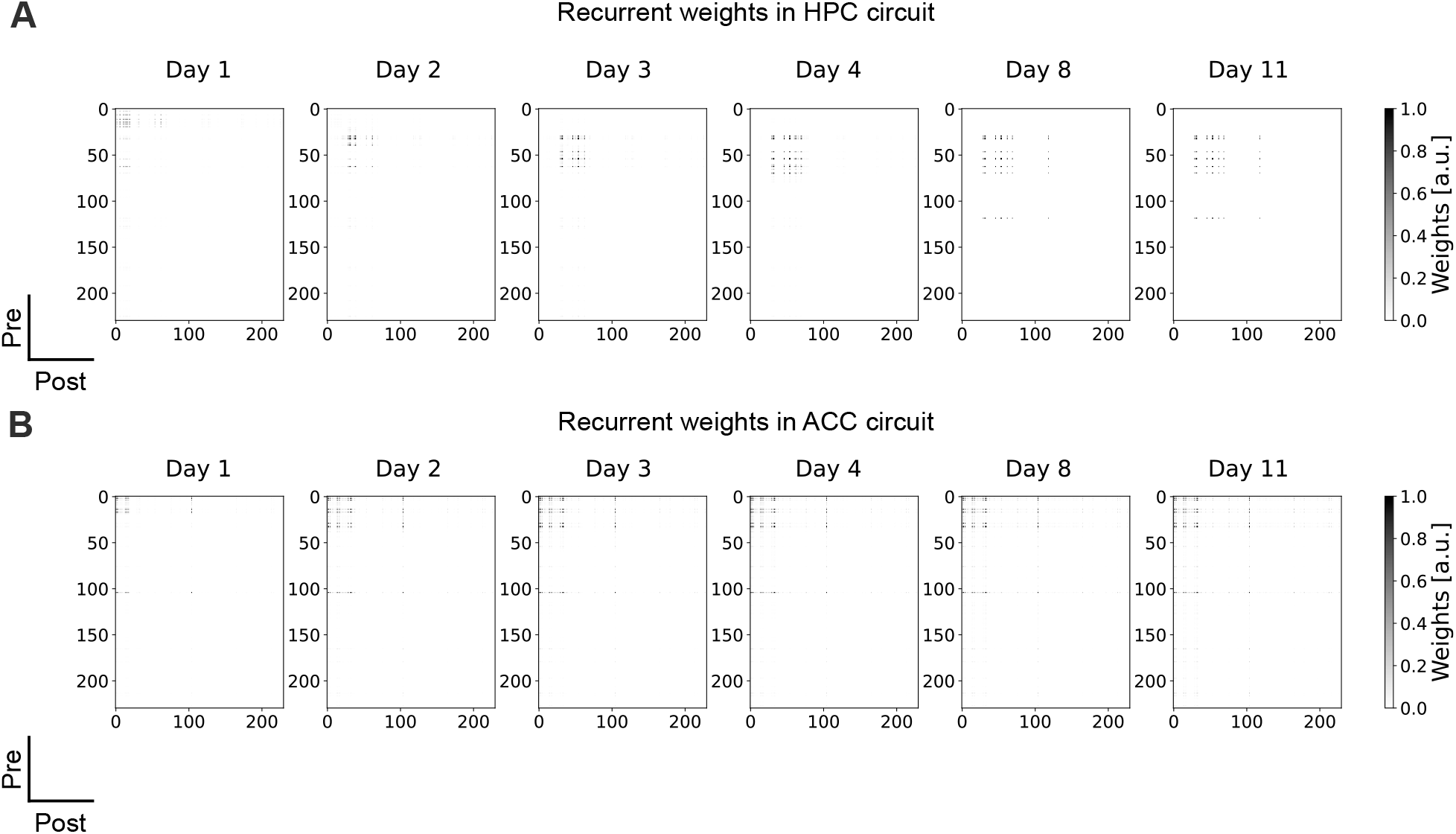
cortical engram drives robust freezing behavior. **(A)** synaptic weights of recurrent connection with-in the HPC network after encoding, and offline reactivation on day 1, 2, 3 8 and 10. **(B)** synaptic weights of recurrent connection in the ACC network after encoding, and offline reactivation on day 1, 2, 3 8 and 10.

**Figure S2:**
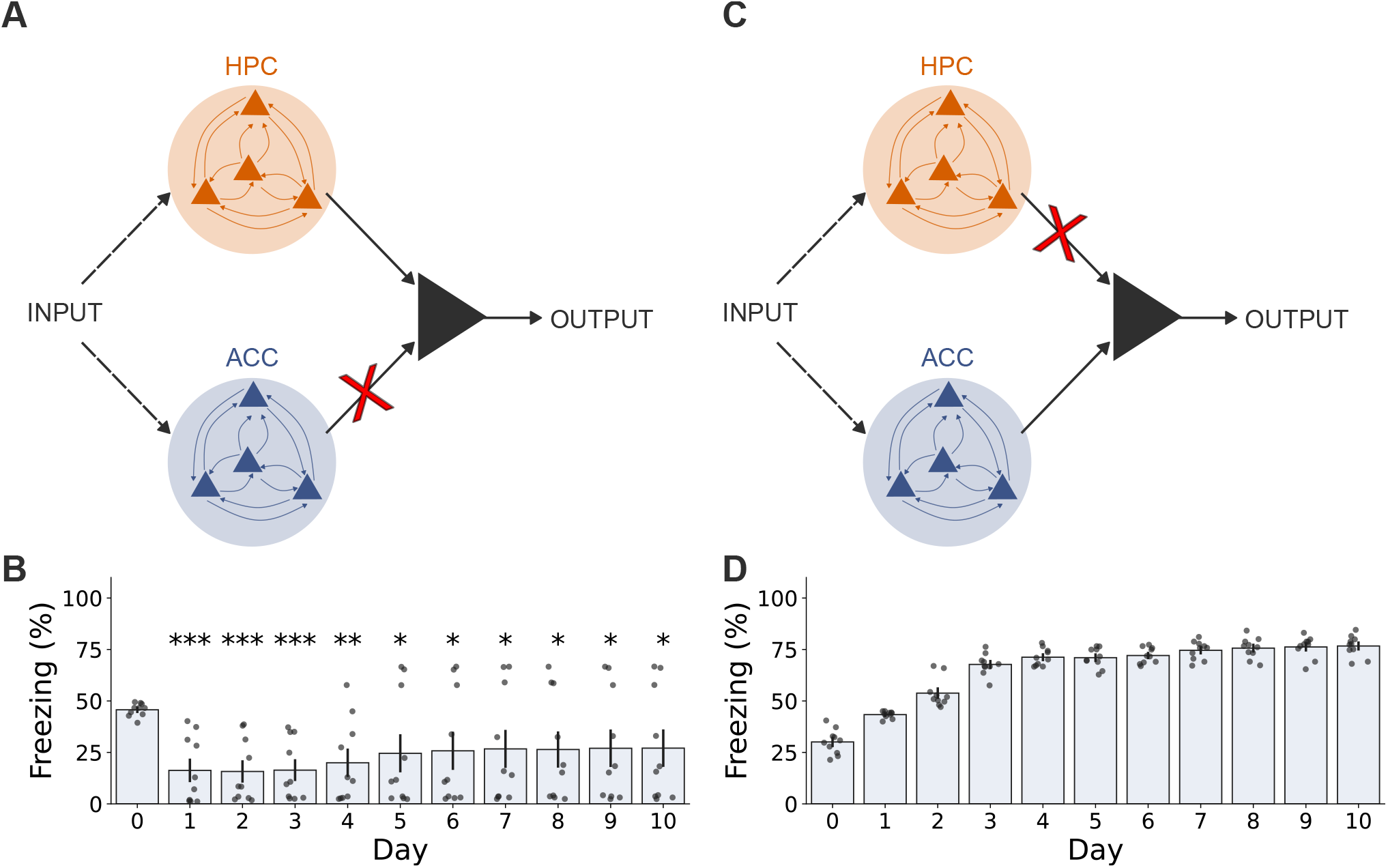
Behavioral consequences of HPC and ACC representational drift. **(A)** Schematic of the HPC-output condition, in which the behavioral output neuron receives plastic input from HPC while ACC-to-output plasticity is disabled.**(B)** Freezing percentage in the HPC-output condition, calculated from output-neuron firing rate (methods) and averaged over 10 stimulus/replay repetitions of each day. Freezing decreases after day 0, indicating that a drifting HPC representation becomes progressively less effective at driving the original behavioral output. **(C)** Schematic of the ACC-output condition, in which the behavioral output neuron receives plastic input from ACC while HPC-to-output plasticity is disabled. **(D)** Freezing percentage in the ACC-output condition, calculated using the same last-10-repetition averaging procedure. Freezing increases across days and approaches a ceiling, indicating that the more stable ACC representation supports persistent and progressively saturated behavioral output. Bars show mean *±* s.e.m across simulations, with individual points representing single simulations.

**Figure S3:**
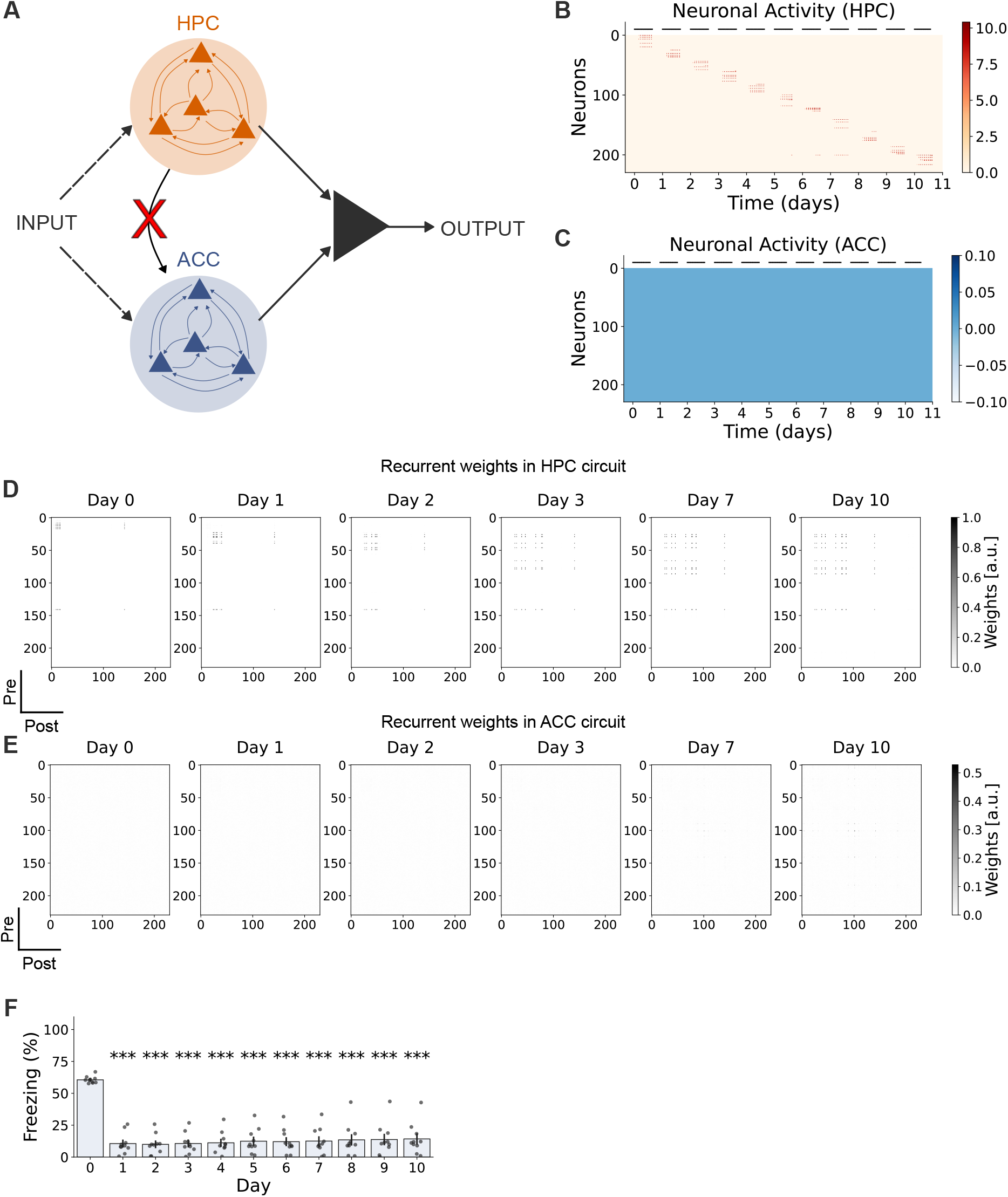
Feedforard projections from HPC to ACC is necesary for engram maturation and stability. **(A)** Schamtic model the model with ablated HPC→ACC feedforward connections. **B** Firing rate of HPC neurons across days. **(C)** Firing of ACC neuron across days. Without the HPC→ACC conenction, the ACC neurons do not mature. **(D)** synaptic weights of recurrent connection with-in the HPC network after encoding, and offline reactivation on day 1, 2, 3 8 and 10. **(E)** synaptic weights of recurrent connection with-in the ACC network after encoding, and offline reactivation on day 1, 2, 3 8 and 10. Syanptic weights do not match without the HPC→ACC connections. **(F)** Without consolidation in ACC, behavioral performance degrades immediately after learning on day 0. Bar plot shows mean and s.e.m of 10 simulations. Individual point represents one simulation.

**Figure S4:**
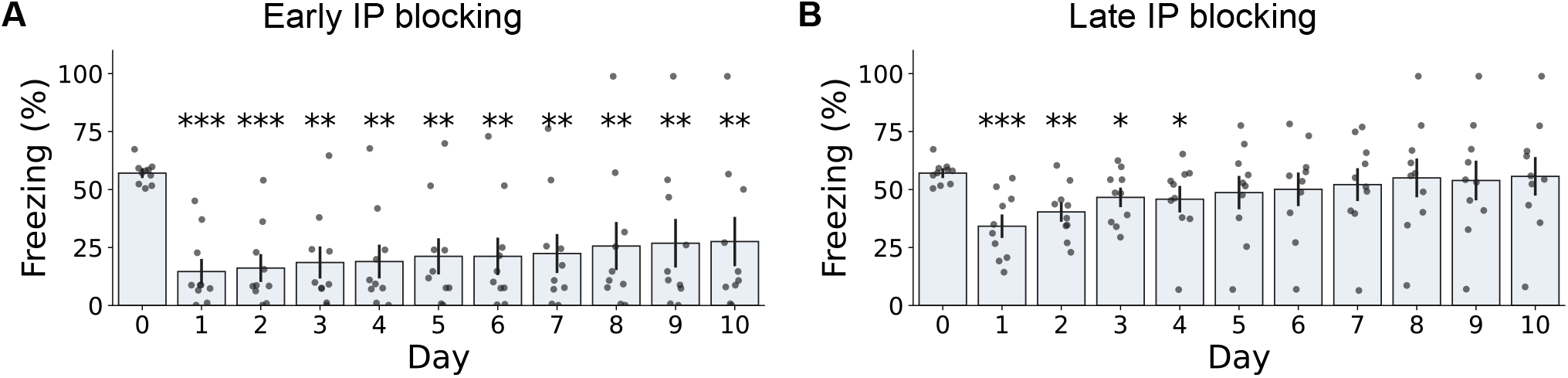
Early IP in ACC is necessary for stable beahvioral. **(A)** Freezing behavior averaged over the 10 presentations of each day. Blocking early (1-2 days after learning) intrinsic plasticity leads to a loss of behavioral output. **(B)** same as **(A)** but in late (4-5 days after learning) IP blocking condition. Late blocking leads to comparable behavioral performance from day 5 onwards until the last day of simulations. Bar plot shows mean and s.e.m of 10 simulations. Individual point represents one simulation.

**Figure S5:**
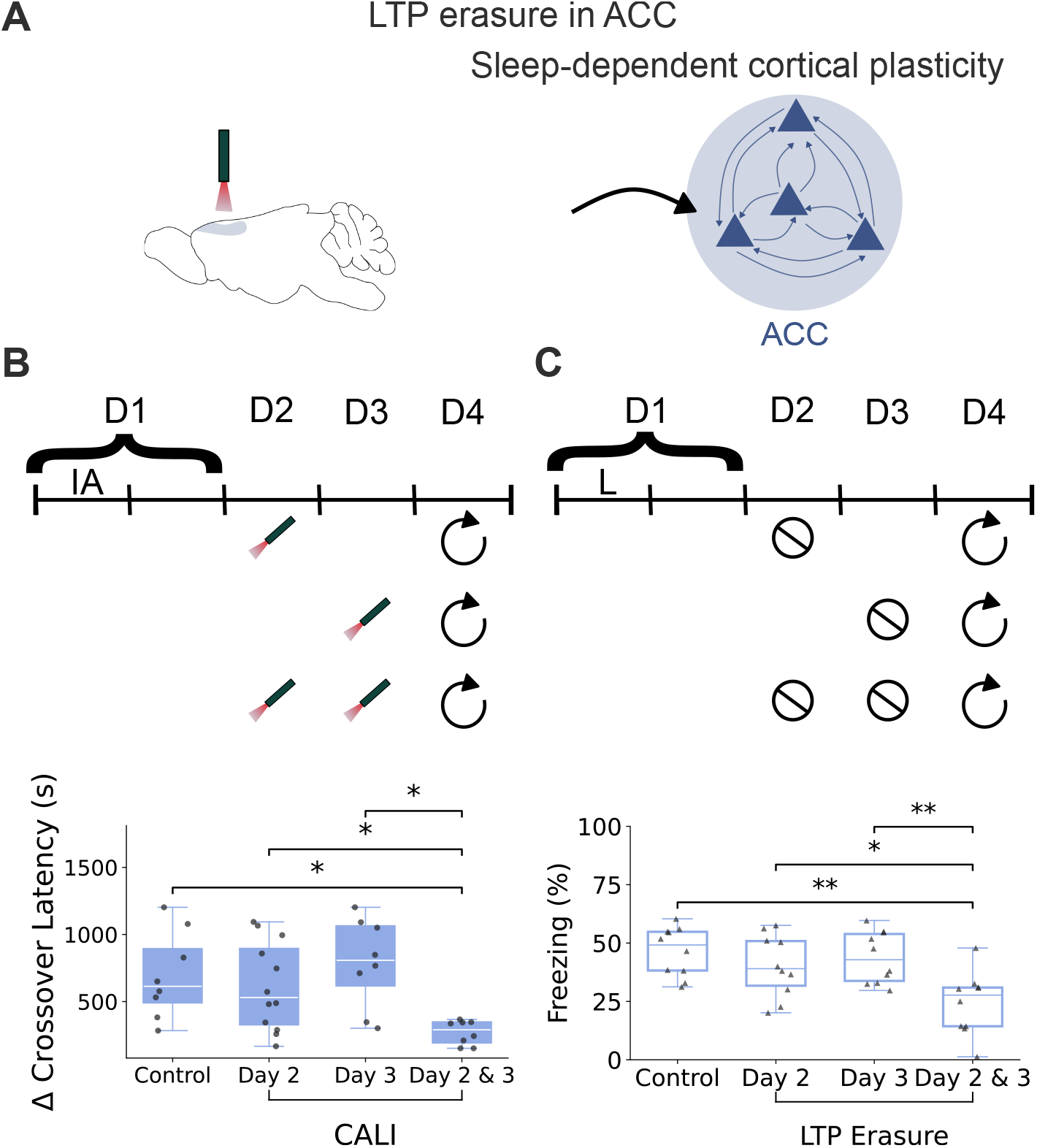
Model reproduces the results on memory restoration by allowing an extra day of plasticity. **(A)** *Left:* Schematic of experimental cortical LTP erasure using CALI in the ACC. *Right:* Schematic of the ACC network in which potentiated recurrent and HPC→ACC feedforward synaptic weights were reset to model LTP erasure (see Methods). **B** *Top left:* Experimental timeline showing ACC LTP erasure using CALI during sleep on day 2, day 3 or both day 2&3. Learning occured on day 1. *Top right:* Corresponding model timeline in which potentiated ACC recurrent and HPC→ACC feedforward weights were reset at the analogous delayed consolidation time point. *Bottom left:* Comparison of memory recall on day 4 across condition. Only CALI Day 2&3 showed memory erasure (Control: N=8, CALI: N(day2) = 12, N(day 3) = 8, N(Day 2&3) = 8, Control vs CALI (Day 2& 3): U=60, corrected p=0.01155 (holm), CALI (Day 2) vs CALI (Day 2 & 3): U=79.5, corrected p=0.01669 (holm), CALI (Day 3) vs CALI (Day 2 & 3): U=58.5, corrected p=0.01249 (holm)). *Bottom right:* In contrast to the experimental result, delayed ACC weight resetting did not substantially reduce freezing in the model (N=10 simulation for each condition, Control vs LTP Erasure Day 2 & 3 : U=94, corrected p=3.2e-3 (holm), LTP Erasure Day 2 vs LTP Erasure Day 2 & 3: U=80, corrected p=2.57e-2 (holm), LTP Erasure Day 3 vs LTP Erasure Day 2 & 3: U=89, corrected p=7.22e-3 (holm)). These results indicate that the current Hebbian plasticity rule does not reproduce the delayed, sleep-dependent requirement for rapid ACC potentiation.

**Figure S6:**
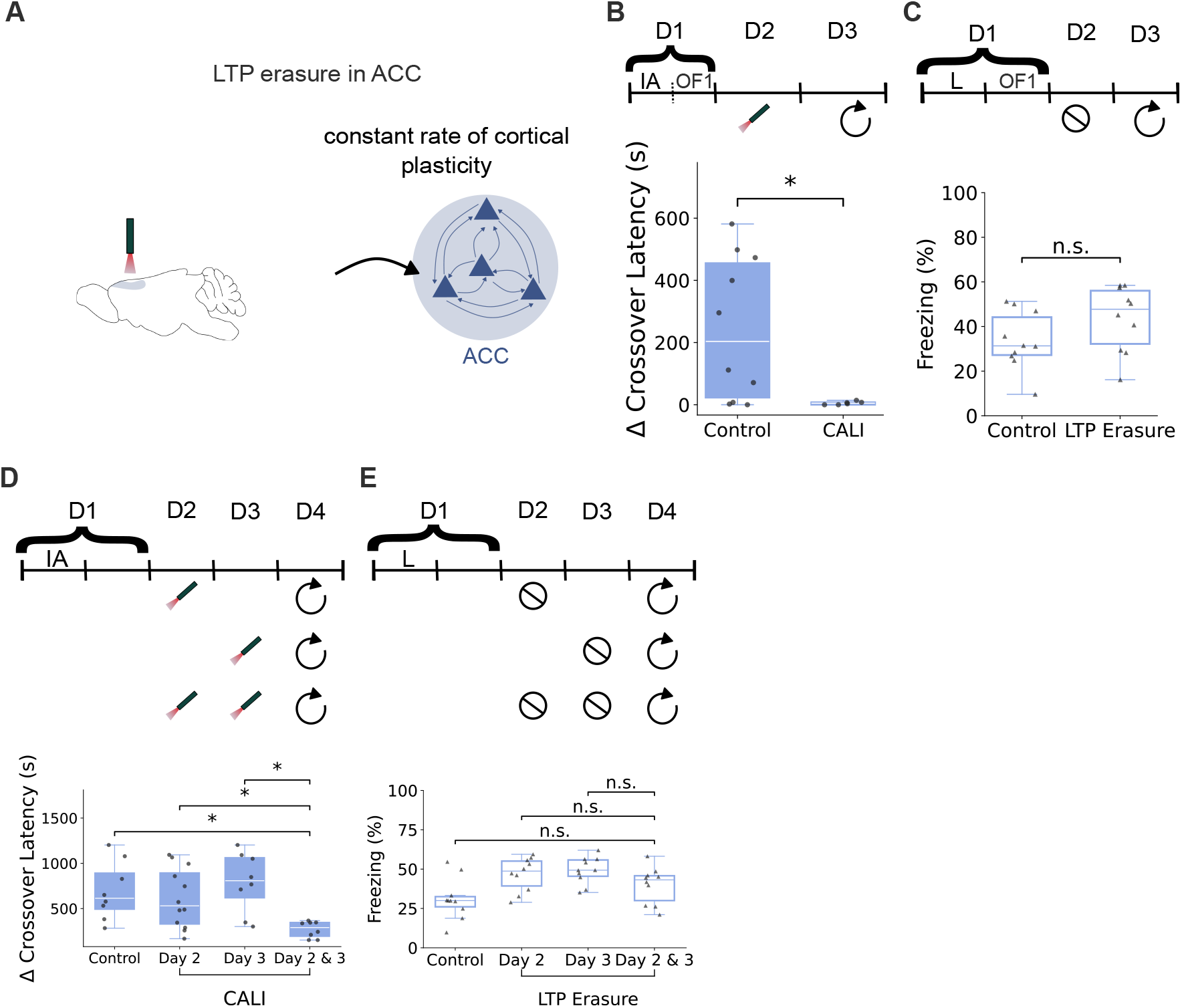
A constant rate of cortical plasticity is unable to explain impact of ACC LTP erasure on behavior. **(A)** *Left:* Schematic of experimental cortical LTP erasure using CALI in the ACC. *Right:* Schematic of the ACC network with a sleep-independent, constant rate of hebbian plasticity, in which potentiated recurrent and HPC→ACC feedforward synaptic weights were reset to model LTP erasure (see Methods). **(B)** *Top:* Experimental timeline showing ACC LTP erasure during sleep on day 2, one day after learning. *Bottom:* Erasure of recently induced ACC LTP during sleep significantly reduced the training-induced increase in crossover latency, indicating impaired inhibitory-avoidance memory (Control: N=10, CALI: N =10, U=49.5, p=4e-2). **(C)***Top:* Corresponding model timeline in which potentiated ACC recurrent and HPC→ACC feedforward weights were reset at the analogous delayed consolidation time point. *Bottom:* In contrast to the experimental result, delayed ACC weight resetting did not substantially reduce freezing in the model (N =10 simulations for each condition, U=29, p=0.73). **(C)** *Top:* Experimental timeline showing ACC LTP erasure using CALI during sleep on day 2, day 3 or both day 2&3. Learning occured on day 1. *Bottom left:* Comparison of memory recall on day 4 across condition. Only CALI Day 2&3 showed memory erasure (Control: N=8, CALI: N(day2) = 12, N(day 3) = 8, N(Day 2&3) = 8, Control vs CALI (Day 2& 3): U=60, corrected p=0.01155 (holm), CALI (Day 2) vs CALI (Day 2 & 3): U=79.5, corrected p=0.01669 (holm), CALI (Day 3) vs CALI (Day 2 & 3): U=58.5, corrected p=0.01249 (holm)). **(E)***Top:* Corresponding model timeline in which potentiated ACC recurrent and HPC→ACC feedforward weights were reset at the analogous delayed consolidation time point. *Bottom:* In contrast to the experimental result, delayed ACC weight resetting did not substantially reduce freezing in the model (N=10 simulation for each condition, Control vs LTP Erasure Day 2 & 3 : U=34, corrected p=0.24 (holm), LTP Erasure Day 2 vs LTP Erasure Day 2 & 3: U=71, corrected p=0.24 (holm), LTP Erasure Day 3 vs LTP Erasure Day 2 & 3: U=75, corrected p=0.19 (holm)). These results indicate that a constant rate of learning for cortical plasticity rule does not reproduce the experimental results and a delayed, sleep-dependent form of plasticity is required for rapid ACC potentiation.

## Notes

### Competing Interest Statement

The authors have declared no competing interest.

